# Three-dimensional correlated random walks for animal movement and habitat selection

**DOI:** 10.1101/2025.11.07.687274

**Authors:** Natasha Klappstein, Théo Michelot, Ron Togunov, Joanna Mills Flemming

**Affiliations:** Department of Mathematics and Statistics, Dalhousie University; Department of Mathematical Sciences, Norwegian University of Science and Technology

**Keywords:** animal movement, habitat selection, step selection function, three-dimensional, 3D, Kent distribution, correlated random walks, biased random walks, telemetry data

## Abstract

Animal movement and habitat selection underpin important ecological phenomena, from individual behaviour to population-level distributions. Despite navigating three-dimensional space, animal movement is typically measured and analysed on a two-dimensional plane, which limits our understanding of animals that swim or fly. Therefore, we propose a step selection function (SSF) capable of quantifying animal movement and habitat selection in three dimensions. We formulate a very general family of three-dimensional correlated random walks, aimed at capturing unique features of three-dimensional data. Using Antarctic petrel data, we show how these SSFs can be used to assess selection for vertically-stratified habitat, account for barriers (e.g., the ground or ocean surface), and model attraction to any number of directional targets. Our modelling framework provides a solid foundation for three-dimensional analyses, which will be crucial to answer ecological questions that would otherwise be ignored in two dimensions.

## 1 Introduction

Animal movement has typically been analysed in two spatial dimensions (e.g., Codling *et al*., 2008; Johnson *et al*., 2008; Avgar *et al*., 2016), while ignoring the vertical dimension (e.g., depth or altitude; Tarroux *et al*., 2016a; Pike & Burman, 2023). Two-dimensional representations of movement may be reasonable for many terrestrial systems, but are clear oversimplifications for marine and volant (i.e., flying) species. For these animals, studying vertical movement is essential to understand their behaviour (Adachi *et al*., 2017; Dunn *et al*., 2024), energetics (Redcliffe *et al*., 2025), resource exploitation (Russo *et al*., 2024), and environmental risks (Péron *et al*., 2017). Therefore, three-dimensional movement analyses hold great promise to improve our knowledge of animal ecology and have been repeatedly identified as a priority for future research (e.g., Tracey *et al*., 2014; Lepczyk *et al*., 2021; Ahmed *et al*., 2022; Lennox *et al*., 2024).

Several existing tracking technologies have the potential to resolve three-dimensional movement, such as GPS-based altitude (Tarroux *et al*., 2016a; Johnston *et al*., 2023), trackers with altimeters (Péron *et al*., 2020) or pressure sensors (Lane *et al*., 2019; Costa *et al*., 2024; Dunn *et al*., 2024), acoustic telemetry systems (Laplanche *et al*., 2015; Udyawer *et al*., 2015; Aspillaga *et al*., 2019; Matley *et al*., 2021), or GPS in conjunction with elevation layers (Tracey *et al*., 2014; Heit *et al*., 2023; Redcliffe *et al*., 2025). Using these data, it has become increasingly common to derive three-dimensional space use (using various methods, e.g., Tracey *et al*., 2014; Ferrarini *et al*., 2018; Aspillaga *et al*., 2019; Demšar & Long, 2019), and these analyses have demonstrated the importance of the vertical dimension in understanding estimating broad-scale animal interactions with habitat (Ferrarini *et al*., 2018), home range size (Tracey *et al*., 2014), and between-individual spatial overlap (Chandler *et al*., 2020; Matantseva & Simonov, 2023). Despite the proliferation of three-dimensional space use estimators, the development of movement models has been comparatively sparse. This limits three-dimensional analyses to only consider large-scale spatial dynamics, while neglecting fine-scale movement, behaviour, and habitat selection.

Although there are several approaches for modelling movement, we focus on developing three-dimensional correlated random walks (CRWs) in discrete time. CRWs capture the speed and directional persistence of movement, and are frequently used in animal movement ecology (Morales *et al*., 2004; Codling *et al*., 2008; Michelot *et al*., 2016). With additional information on the animal’s vertical position, we can specify a CRW compatible with three-dimensional step lengths and directional changes (Benhamou, 2019). Recent methodological work has just begun to recognise the utility of three-dimensional CRWs for animal tracking data, including both positional data (Benhamou, 2019; Ahmed *et al*., 2021; Pike & Burman, 2023) and high-resolution biologging data (Benhamou, 2023). However, adding the third dimension creates unique challenges for movement modelling, many of which have not been solved. One important challenge arises from the inequality of scales between dimensions, due to additional energetic costs or physical constraints that limit movement in the vertical dimension (as identified in Pike & Burman, 2023). It can be difficult to reconcile these scales in a single movement model, and may explain why many (quasi-3D) studies rely on treating horizontal and vertical data disparately from one another (e.g., Bestley *et al*., 2015; McClintock *et al*., 2013; Heit *et al*., 2023; Pritchard *et al*., 2025). Previous papers on three-dimensional CRWs have developed important theory, but there has been little emphasis on parameter estimation, resolving the biological discrepancy between the horizontal and vertical scales, or extension to more complex ecological contexts.

Three-dimensional CRWs can serve as the basis for more representative analyses of animal behaviour and space use. Here, we propose a step selection function (SSF), capable of resolving three-dimensional movement and habitat selection. An SSF defines the likelihood of a step ending at location ***s***_*t*+1_ given the previous locations ***s***_1:(*t*−1)_ = {***s***_1_, ***s***_2_, …, ***s***_*t*−1_} as,

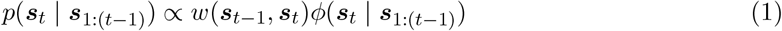

where *w* is a habitat selection function that describes the habitat preferences of the animal and *ϕ* is a movement kernel. In two dimensions, SSFs have been widely used to analyse fine-scale animal movement processes, such as responses to linear features (Prokopenko *et al*., 2017), spatial memory (Kim *et al*., 2024), inter-individual interactions (Potts & Börger, 2022), and behaviour (Nicosia *et al*., 2017; Klappstein *et al*., 2023). Since SSFs capture the spatial mechanisms of animal movement, they can be used to estimate large-scale patterns of space use (Signer *et al*., 2017; Potts & Börger, 2022; Signer *et al*., 2024).

In this paper, we propose specifying *ϕ* as a three-dimensional CRW, which allows insights into vertically-dependent behaviour and selection for three-dimensional habitat variables in *w*. To do so, our main objectives are to: i) define appropriate movement metrics to quantify three-dimensional movement, ii) formulate a general class of three-dimensional CRWs capable of resolving observed movement patterns, and iii) integrate these CRWs into the SSF framework. Using theory, simulations, and an example of Antarctic petrels (*Thalassoica antarctica*; data from Tarroux *et al*., 2016a,c), we show how our method can be used to capture inequality between the horizontal and vertical dimensions, account for three-dimensional barriers, assess selection for vertically-stratified habitat, and model attraction to any number of directional targets.

## 2 Quantifying three-dimensional movement

Two-dimensional movement speed and directional persistence are typically measured via step lengths and turning angles (Codling *et al*., 2008). Here, following Benhamou (2019, 2023), we define appropriate metrics to quantify speed and directional persistence in three dimensions. Note, throughout this paper, we will be referring to both Cartesian and spherical coordinate systems extensively. Further, the unit sphere will be an important concept used to define three-dimensional angular variables. If these concepts are unfamiliar, see Appendix A.1 for a brief refresher.

Suppose we have animal locations ***s***_*t*_ composed of three-dimensional Cartesian coordinates (easting, northing, altitude/depth), ***s***_*t*_ = {*x*_*t*_, *y*_*t*_, *z*_*t*_}, collected at regularly-spaced times *t* ∈ {1, 2, …, *T*}. Note that we consider the animal’s position in projected space, and assume that all three dimensions are measured in the same unit (e.g., metres). Movement can be described as a sequence of three-dimensional steps Δ***s***_*t*_ = ***s***_*t*_ − ***s***_*t*−1_ = {Δ*x*_*t*_, Δ*y*_*t*_, Δ*z*_*t*_}, as illustrated in Figure 1a. As in two dimensions, the step length *l*_*t*_ ∈ [0, ∞) is defined as the Euclidean distance between two successive positions, 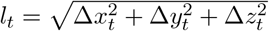. The direction of movement can be summarised by two angles: the horizontal (i.e., azimuthal) bearing *θ*_*t*_ and the vertical (i.e., polar) bearing *α*_*t*_, defined as

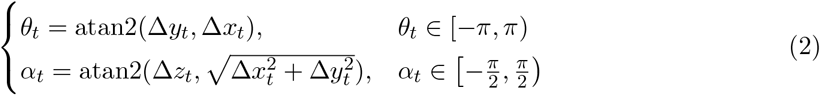

where atan2 is the two-argument arctangent function. The horizontal bearing represents the cardinal direction of movement (East = 0; West = *±π*) and the vertical bearing indicates whether the animal moved up or down between locations. Like Benhamou (2019), we define vertical bearing relative to the horizontal plane (where level movement = 0; Figure 1b), as animals may favour level movements. The two bearing angles *θ*_*t*_ and *α*_*t*_ are the spherical coordinates of a point on the three-dimensional unit sphere (of radius 1). This point has Cartesian coordinates ***v***_*t*_ = {cos(*α*_*t*_) cos(*θ*_*t*_), cos(*α*_*t*_) sin(*θ*_*t*_), sin(*α*_*t*_)}, and can also be calculated as ***v***_*t*_ = (Δ***s***_*t*_)*/l*_*t*_ (i.e, the step vector scaled to length 1; Figure 1b). Therefore, the coordinates {*θ*_*t*_, *α*_*t*_} and ***v***_*t*_ can be used interchangeably to describe the direction of movement at time *t*.

**Figure 1:**
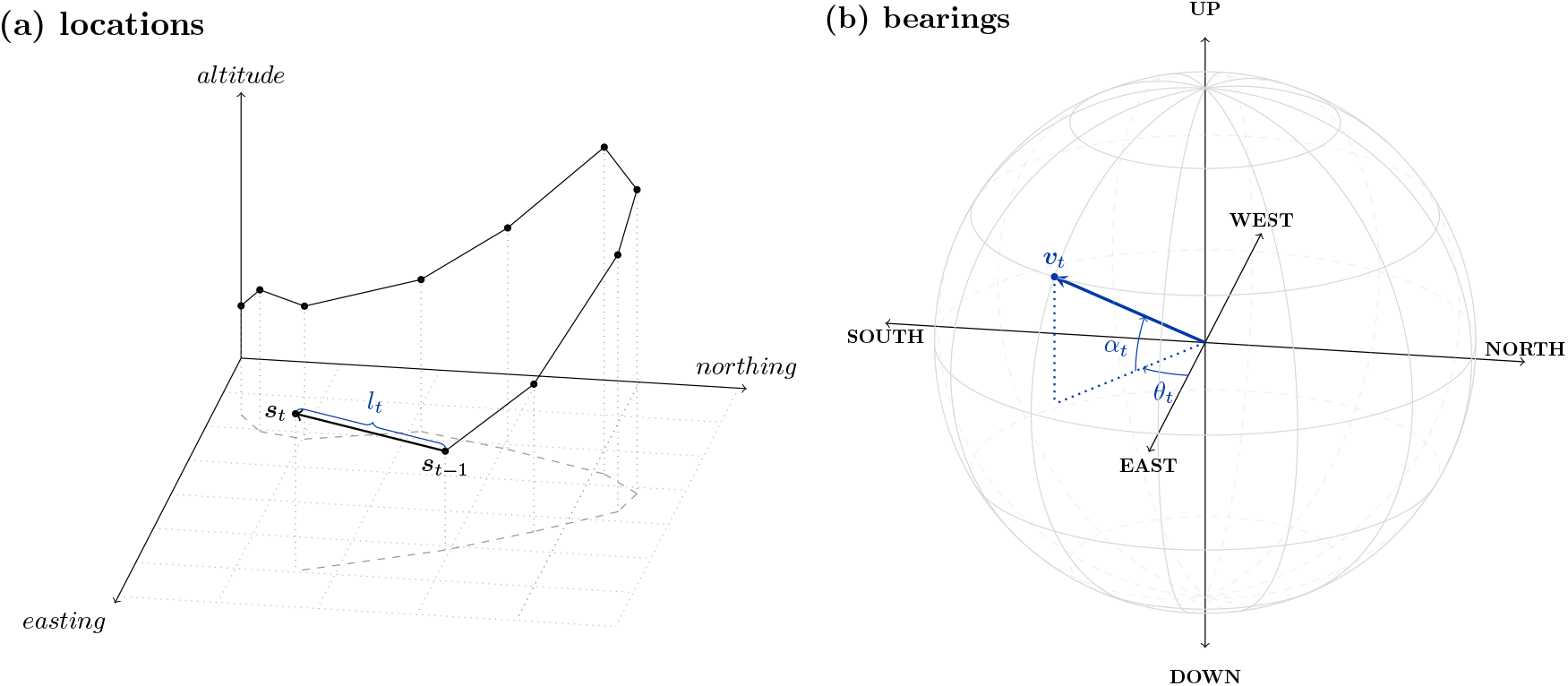
Hypothetical example of three-dimensional movement data. (a) A track ending at location **s**_t_, where the step length l_t_ is the Euclidean distance between successive locations. (b) A bearing on the unit sphere representing the direction of travel. The bearing in (b) can be described by its spherical coordinates {θ_t_, α_t_} or Cartesian coordinates **v**_t_.

To capture directional persistence, we are interested in the change in bearings between successive steps. Naively, we might consider taking simple differences between successive bearings, defining the vertical turning angle as 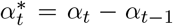 and the horizontal turning angle as 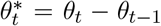. Although 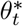 is sufficient to capture directional persistence in two dimensions, 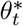 and 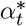 do not adequately describe a directional change in three dimensions. That is, a given combination of 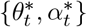 could correspond to very different changes in direction, and cannot be readily interpreted (see Appendix A.2 for an illustrative example). Rather than using 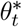 and 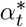, a valid representation of a three-dimensional change in direction is a geodesic (i.e., the shortest arc connecting points on a sphere; Benhamou, 2019). If we have two successive bearings ***v***_*t*−1_ and ***v***_*t*_ on the surface of a unit sphere, we are interested in the geodesic that connects them (Figure 2a). The geodesic has length

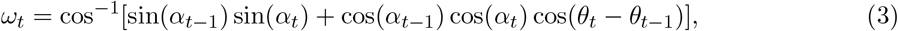

where *ω*_*t*_ ∈ [0, *π*]. The arc length describes the magnitude of change, where small values indicate a high level of directional persistence and values close to *π* indicate a large reversal in direction. To properly specify the geodesic, we also define a direction (relative to some reference) as

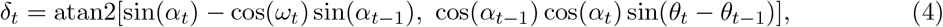

where *δ*_*t*_ ∈ (−*π, π*] (Benhamou, 2019, 2023). The reference (i.e., where *δ* = 0) is based on great circle geometry. Consider we have two great circles that both intersect the bearing ***v***_*t*−1_: i) the first circle goes through the poles of the unit sphere, and ii) the second circle is orthogonal to the first. Then, *δ*_*t*_ is the angle between the second great circle and the geodesic (Figure 2a). When the change in direction is along this orthogonal great circle, *δ*_*t*_ = 0 or *δ*_*t*_ = *π*. Conversely, if the change in direction is along the first great circle, *δ*_*t*_ = *±π/*2. The biological interpretation of *δ*_*t*_ is complicated because it depends on the previous direction of movement {*θ*_*t*−1_, *α*_*t*−1_}, but we can develop intuition in some special cases. Importantly, when the previous vertical bearing *α*_*t*−1_ = 0 (i.e., corresponding to level movement), the orthogonal great circle is the equator of the unit sphere. In that case, a purely horizontal change in direction corresponds to *δ* = 0 (change to the left) or *δ* = *π* (change to the right). We consider that this is an important case in animal movement, as many species may tend to take steps with relatively level bearings. Note that we assume that an animal’s posture remains constant, and do not consider the bank angle as in Benhamou (2023).

**Figure 2:**
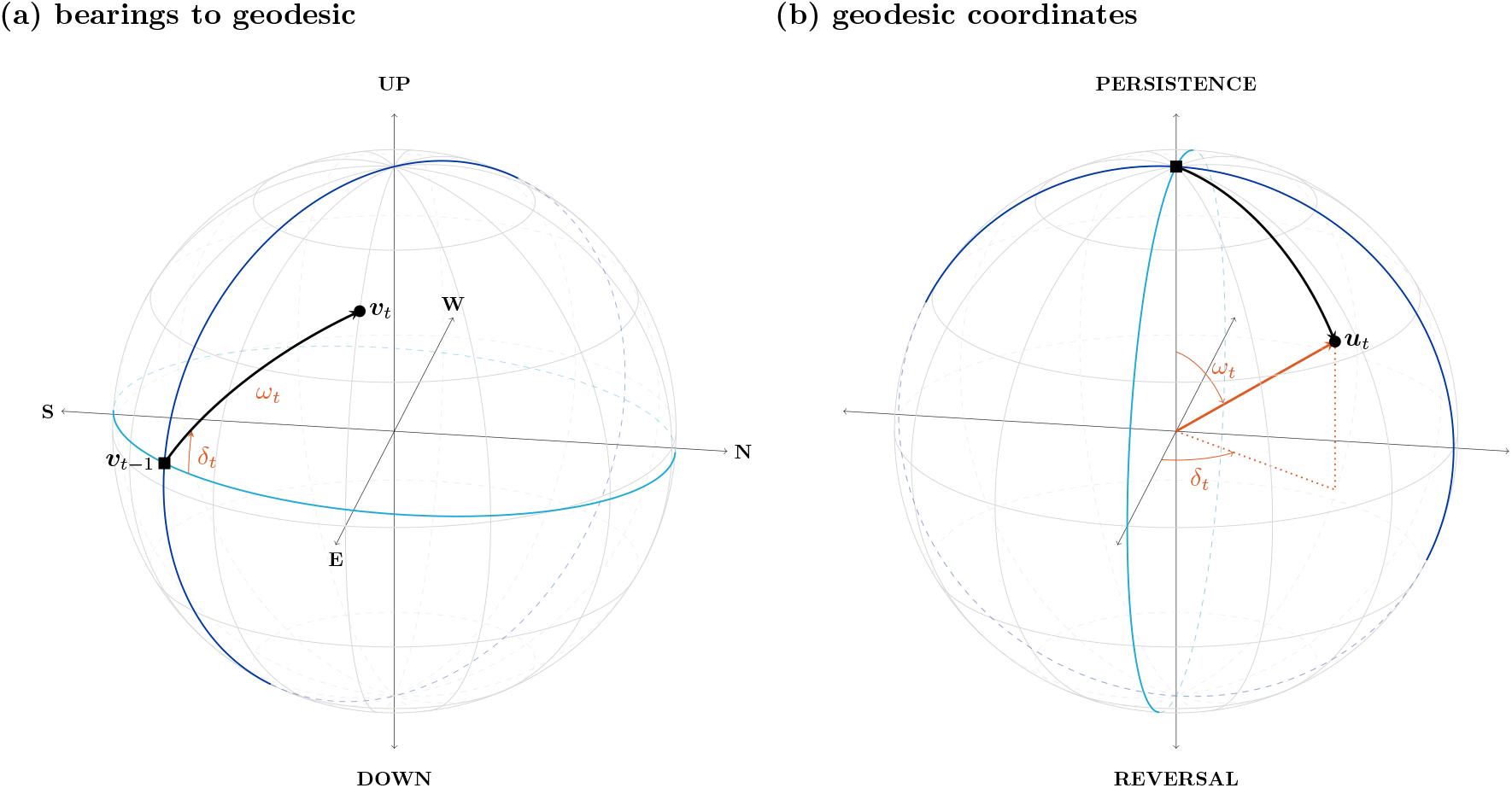
Illustration of how to derive geodesics from successive bearings. (a) The geodesic (black line) is the shortest arc connecting two bearings **v**_t−1_ and **v**_t_. Both blue great circles pass through **v**_t−1_: the dark blue great circle goes through the poles, and the light blue great circle is orthogonal to the dark blue circle. ω_t_ is the arc length of the geodesic, and δ_t_ is the angle between the light blue great circle and the geodesic. (b) Demonstration of the geodesic as a point on the unit sphere. The geodesic can be described in terms of its Cartesian coordinates **u**_t_ or spherical coordinates: ω_t_ is the polar/vertical angle measured from the vertical axis, and δ_t_ is the horizontal angle. The blue great circles are the same in the two panels, such that (b) can be viewed as a rotation of (a), where the previous bearing **v**_t−1_ is at the “North Pole”, and where the axes are aligned with the great circles. Note that the unit sphere in (a) represents bearings and therefore its axes correspond to direction of travel, whereas (b) represents directional changes (where “up” is directional persistence).

Taken together, *ω*_*t*_ and *δ*_*t*_ define a three-dimensional directional change. Like bearings, these two angles can be viewed as the spherical coordinates of a point on a unit sphere (Figure 2b), and this point can equivalently be written in terms of its Cartesian coordinates

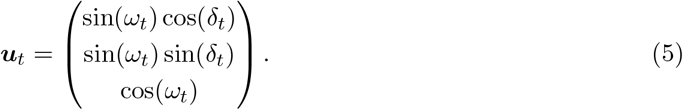

Note that this transformation is different than that of bearings as we use are measuring the vertical angle (i.e., *ω*_*t*_) relative to the vertical axis, rather than the horizontal plane (as we do for *α*_*t*_). Therefore, 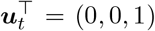 corresponds to directional persistence (i.e., *ω*_*t*_ = 0). Although angular variables *ω*_*t*_ and *δ*_*t*_ are an intuitive description of the geodesic, statistical models are often written in terms of ***u***_*t*_.

## 3 Three-dimensional correlated random walks

In discrete time, two-dimensional CRWs are commonly defined as probability distributions of polar coordinates (i.e., step length and turning angle), and this framework can be extended to accommodate three-dimensional movement metrics (i.e., step lengths and geodesics; Benhamou, 2019; Ahmed *et al*., 2021; Benhamou, 2023). Three-dimensional step lengths can be modelled with any appropriate probability distribution with support *l >* 0 and a sensible choice is the gamma distribution (Morales *et al*., 2004; Langrock *et al*., 2012). When parametrised by its shape *a* and scale *b*, it has the following density

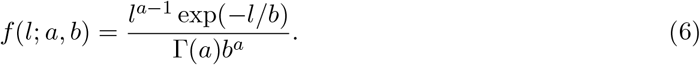

To form a CRW, we combine Equation 6 with a distribution of directional changes. Next, we propose three angular models for three-dimensional data.

### 3.1 Isotropic CRW

If we assume isotropy (i.e., that any geodesic orientation *δ*_*t*_ is equally likely) then directional changes can be modelled with a spherical von Mises Fisher (vMF) distribution of the unit vector of geodesics ***u***_*t*_ (e.g., Equation 5),

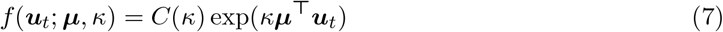

where ***µ*** is a unit vector representing the mean direction, *κ >* 0 is the angular concentration parameter, and *C*(*κ*) is a normalisation constant (Mardia & Jupp, 2009). The vMF can be of any positive dimension *p*; when *p* = 2 the vMF reduces to a von Mises distribution on a circle, which is typically used in two-dimensional movement studies. If we assume that animals exhibit directional persistence, we can set ***µ***^⊤^ = (0, 0, 1) and then the density can be written,

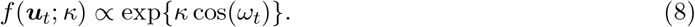

The vMF is an isotropic distribution (i.e., it is symmetric around its mean). In the context of directional changes, this assumes that animals change direction with equal magnitude in both the horizontal and vertical planes of movement, which may not always be realistic (as recognized by Pike & Burman, 2023). Therefore, in the next two sections, we propose two extensions of the isotropic CRW to solve this problem: i) a CRW with a Kent distribution of geodesics, and ii) a biased correlated random walk (BCRW).

### 3.2 Kent CRW

The Kent distribution is a generalisation of the vMF distribution that relaxes the assumption of isotropy (Kent, 1982). The Kent distribution is analogous to a bivariate normal distribution on a sphere, and specifies an elliptical density,

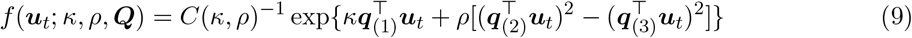

where *κ* ≥ 0 is the concentration, *ρ* is a parameter than controls the ovalness of the ellipse (where 0 ≤ *ρ < κ/*2), and *C*(*κ, ρ*) is the normalisation constant. The location of the ellipse is controlled by the orthogonal matrix ***Q*** = {***q***_(1)_, ***q***_(2)_, ***q***_(3)_}, whose columns describe the mean direction, major axis, and minor axis (respectively). When *ρ* = 0, the Kent distribution reduces to a symmetric vMF distribution (Equation 7).

We propose a simplification of the Kent distribution based on biological assumptions, to reduce the number of parameters to estimate and facilitate its use in an SSF. We can fix the elements of ***Q*** to assume that the mean is 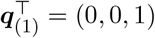, the major axis is 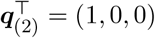, and the minor axis is 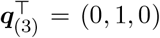. These assumptions are meant to capture two anticipated properties of animal movement: i) directional persistence (via the mean vector), and, ii) higher variability of horizontal bearings, compared to vertical bearings (via the major/minor axes). Under this frame of reference (and ignoring the constant), the density of the Kent distribution simplifies to

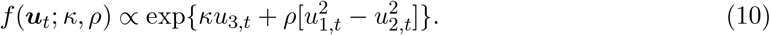

where *u*_1,*t*_, *u*_2,*t*_, *u*_3,*t*_ are the entries of ***u***_*t*_ (Kent, 1982). Within this simplified formulation, it is possible to allow *κ* and *ρ* to be negative: a negative *κ* is equivalent to setting the mean to 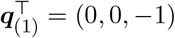 (representing directional reversal), and a negative value of *ρ* swaps the major and minor axes of the distribution (allowing for more vertical movement than horizontal movement). Combining Equations 10 and 5, the density can be written in terms of spherical coordinates as

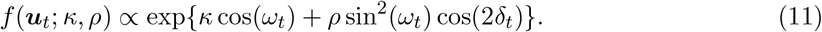

The ovalness parameter is constrained to be less than *κ/*2, which limits the capacity of this model to capture very stretched distributions, i.e., with strong anisotropy in movement direction. Therefore, next, we also propose an alternative solution based on a BCRW that makes no assumptions about the degree of anisotropy.

### 3.3 Biased CRW

In a BCRW, we suppose that an animal’s direction is a compromise between directional persistence and bias towards target directions (Codling *et al*., 2008; Benhamou, 2014). BCRWs have been explored extensively in two dimensions, (including for SSFs; Duchesne *et al*., 2015; Nicosia *et al*., 2017). Here we define a three-dimensional version, based on the “consensus” model of Duchesne *et al*. (2015) and Rivest *et al*. (2016). We start by describing the general BCRW in 3D, and later we propose a useful special case to account for anisotropy.

Consider we want to model an animal’s direction at time *t* as a compromise between *K* directional targets. We denote as 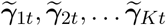 the unit vectors of the target bearings, one of which is the animal’s previous bearing to capture directional persistence (e.g., 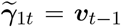). We aim to model the animal’s bearing ***v***_*t*_ with a vMF distribution,

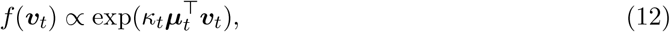

where ***µ***_*t*_ is the mean direction and *κ*_*t*_ is the concentration parameter, both of which depend on the directional targets. Specifically, both parameters can be defined in terms of the weighted sum 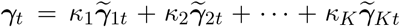, where each unit vector 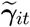 is weighted by the relative attractiveness of target *i* (measured by *κ*_*i*_). Then, the mean direction is defined as ***µ***_*t*_ = ***γ***_*t*_*/*∥***γ***_*t*_∥ (i.e., the direction of ***γ***_*t*_) and *κ*_*t*_ = ∥***γ***_*t*_∥ as the length of ***γ***_*t*_. In this model, the bearing of the next movement step of the animal is closely aligned with 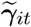 if *κ*_*i*_ is large, and the turning angles have higher concentration (lower variance) when the directional targets are all in the same direction. Therefore, the target weights {*κ*_1_, *κ*_2_, …, *κ*_*K*_} are not time-varying, but the overall concentration *κ*_*t*_ is because it depends on the agreement between targets at time *t*. Substituting these definitions of *κ*_*t*_ and ***µ***_*t*_ into Equation 12, the density can be written as

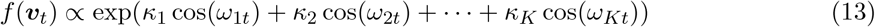

where *ω*_*it*_ is the arc length of the geodesic between ***v***_*t*_ and 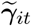 (full calculations in Appendix B.2). Equation 13 is very flexible and can incorporate any directional targets; without attraction to the previous heading ***v***_*t*−1_, this model reduces to a biased random walk (Ahmed *et al*., 2021).

To account for anisotropy in directional changes, we wish to consider the animal’s bearing at time *t* as compromise between persistence and level movement. By analogy with the BCRW developed above, we model ***v***_*t*_ using Equation 13, based on two geodesic lengths *ω*_1*t*_ and *ω*_2*t*_. Here, *ω*_1*t*_ is the length of the geodesic connecting ***v***_*t*_ and ***v***_*t*−1_ (to capture persistence, as in the isotropic case). To capture the tendency to take level steps with a small vertical bearing, we define *ω*_2*t*_ as the arc length between ***v***_*t*_ and the closest bearing with a vertical bearing of 0 (i.e., the unit vector with spherical coordinates {*θ*_*t*_, 0}). Since the vertical bearing *α*_*t*_ is measured relative to the horizontal plane, by definition, *ω*_2*t*_ = |*α*_*t*_|, such that the model can be written

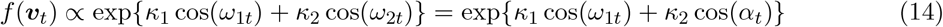

Compared to the Kent CRW, the BCRW may have greater flexibility to capture extreme ovalness, since the direction of travel can be heavily weighted towards level movement via *κ*_2_.

## 4 Three-dimensional step selection functions

The previous sections focused on formulating sensible CRWs to describe animal movement. Here, we explain how to combine a three-dimensional (B)CRW and habitat selection into an SSF, and show that this model can be implemented using convenient methods and software.

### 4.1 Model formulation

The SSF likelihood (expanded from Equation 1) of a step ending at ***s***_*t*_ is

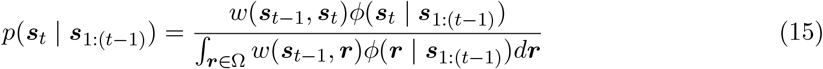

where *w* is the habitat selection function, *ϕ* is the movement kernel (typically modelling a CRW), and Ω is the spatial domain of integration (Forester *et al*., 2009; Avgar *et al*., 2016; Michelot *et al*., 2024). Equation 15 is a probability density function over two- or three-dimensional space, representing where the animal is likely to go at time *t*. Typically, both *w* and *ϕ* are specified as log-linear models: 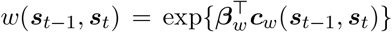 and 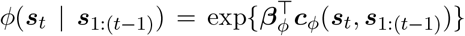. Therefore, the movement kernel and habitat selection function can be factorised to get a single linear predictor,

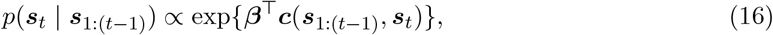

where ***β***^⊤^ = (***β***_*w*_, ***β***_*ϕ*_) and ***c***(***s***_1:(*t*−1)_, ***s***_*t*_)^⊤^ = (***c***_*w*_(***s***_*t*−1_, ***s***_*t*_), ***c***_*ϕ*_(***s***_*t*_ | ***s***_1:(*t*−1)_)).

Here, we describe how to derive the variables needed in ***c***_*ϕ*_(***s***_*t*_, ***s***_1:(*t*−1)_) to model the CRWs in Section 3. The general idea is to write the likelihood of a point ***s***_*t*_ under a given CRW and match this expression with Equation 16. Importantly, the likelihood *p*(***s***_*t*_ | ***s***_1:(*t*−1)_) is a distribution over three-dimensional Cartesian space, and so we need a change of coordinates for any distributions of spherical variables (Rhodes *et al*., 2005; Schlägel & Lewis, 2016; Michelot *et al*., 2024). We start with the isotropic CRW, where we model step lengths *l*_*t*_ with a gamma distribution and directional changes ***u***_*t*_ with a vMF (where *l*_*t*_ and ***u***_*t*_ are independent). The expression for the vMF distribution, *f* (***u***_*t*_) ∝ exp{*κ* cos(*ω*_*t*_)} (Equation 8), is already a density over Cartesian space, but the step length distribution is not a spatial density. A transformation is required because there is more spatial volume far from the start point ***s***_*t*−1_ than close to it (Rhodes *et al*., 2005). Specifically, the volume of a spherical shell is proportional to the square of its radius, so the spatial distribution implied by a given distribution *f* (*l*_*t*_) of step lengths is proportional to 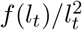. We explain this in more detail in Appendix C.1, and also show how to obtain the transformation via a change-of-variable method. Then, combining the gamma (Equation 6) and the vMF distributions, the isotropic CRW can be written

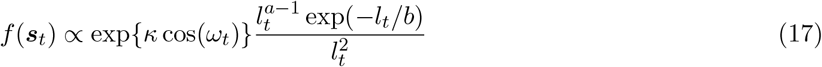

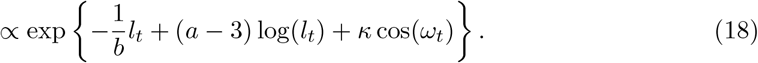

By analogy with Equation 16, this corresponds to an SSF with the variables ***c***_*ϕ*_(***s***_*t*_, ***s***_1:(*t*−1_) = {*l*_*t*_, log(*l*_*t*_), cos(*ω*_*t*_)} and parameters 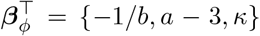. We use a similar reasoning for the anisotropic models, noting that the variables and coefficients remain the same for the gamma distribution of *l*_*t*_. We find that the Kent distribution has the angular variables {cos(*ω*_*t*_), sin^2^(*ω*_*t*_) cos(2*δ*_*t*_)} with corresponding parameters {*κ, ρ*}. The BCRW takes the same form as Equation 13 and can include any number of targets via the covariates {cos(*ω*_1*t*_), cos(*ω*_2*t*_), … cos(*ω*_*Kt*_)} whose coefficients correspond to the relative attractiveness of each target. Note that one of the arc lengths will typically correspond to the arc length between successive bearings and is therefore an extension of Equation 18 with additional target directions.

By formulating our model as an SSF, these CRWs can be combined with any number of (potentially three-dimensional) environmental variables to jointly estimate movement with habitat selection via exp 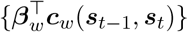. The habitat selection function can include any covariates that may affect an animal’s movement decisions, such as foraging resources or proxies of risk (e.g., Prokopenko *et al*., 2017; Anton *et al*., 2020; Nisi *et al*., 2022). The coefficients ***β***_*w*_ measure preference (*β >* 0) and avoidance (*β <* 0) of these spatial features, and are typically interpreted in terms of relative selection strength (Avgar *et al*., 2017; Fieberg *et al*., 2021). Our approach can utilise three-dimensional variables to assess selection for vertically-stratified resources or features. Further, since the SSF is a spatially-explicit model, it is a natural framework to account for movement barriers, which may be important to consider in three dimensions (e.g., the ocean’s surface and floor). We illustrate these opportunities with an example of petrels in Section 5.

### 4.2 Parameter estimation

The full track likelihood is the product of all step likelihoods (Equation 15),

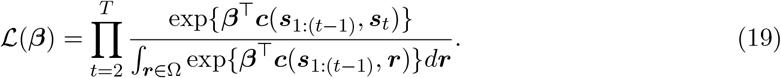

The denominator is a normalising constant needed to ensure that the model is a valid probability density function, which is continuous over three-dimensional space (i.e., representing an infinite number of locations the animal could choose from). However, we have no analytical way to evaluate this complex integral. The typical approach is to approximate the function with numerical integration methods or a discrete-choice approximation (although these methods are equivalent; Michelot *et al*., 2024). To approximate the step likelihood at each time *t*, each observed location is matched with *N* random points sampled from some distribution *h*. This leads to a stratum with a single observed location matched with {***r***_1*t*_, ***r***_2*t*_, … ***r***_*Nt*_} random points and the following approximate likelihood,

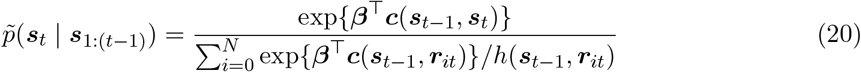

where ***r***_0*t*_ is the observed location. Here, *h* is a spatial distribution to approximate the three-dimensional integral, and the choice of *h* can affect the accuracy and precision of parameter estimates (Michelot *et al*., 2024). In the three-dimensional case, we expect that efficient sampling may be even more important, as adding even a single dimension greatly increases the number of points needed to approximate a given density (i.e., the “curse of dimensionality”). See Appendix C.2 for a more detailed description of uniform and importance sampling.

A convenient approach to estimate the parameters for the model in Equation 20 is conditional logistic regression (CLR), which is available in existing software such as the `survival, amt`, and `mgcv` packages in R (Wood, 2017; Signer *et al*., 2019; Therneau, 2022). Equation 20 is almost equivalent to a CLR model, except for the function *h* in the denominator, and parameters estimated by CLR much be corrected to reflect our choice of *h* (Forester *et al*., 2009; Avgar *et al*., 2016). If *h* is spatially uniform, then this correction is not needed, but this is typically inefficient (Michelot *et al*., 2024). In our analyses (Section 5), we focus on using importance sampling where random points are sampled with a gamma distribution of step lengths and uniform bearings (see Appendix C.3 for detailed parameter corrections). We conducted simulations to test our proposed implementation under various forms of *h* and choices of *N* (Appendix D.2). We showed that our SSF estimators are unbiased for all three CRWs, when using importance sampling. It is important to note that this implementation is a simple extension of the standard SSF workflow (outlined in Klappstein *et al*., 2024): i) sample three-dimensional integration points, ii) evaluate movement and habitat variables of interest, iii) fit SSF with CLR software, and iv) interpret parameters (with corrections, if necessary).

## 5 Illustration

To illustrate our method, we analysed three-dimensional tracking data from Antarctic petrels, obtained from the MoveBank Data Repository (Tarroux *et al*., 2016a,c). We conducted two main analyses: i) a preliminary comparison of all movement kernel specifications in Section 3, and ii) an exploration of how petrels use wind to their energetic advantage (based on wind support, following Tarroux *et al*., 2016a). Analysis (ii) also explores how inferences differ between models with and without vertical information about habitat (i.e., wind). Note these analyses were designed to demonstrate our methodology, rather than to thoroughly investigate petrel ecology.

### 5.1 Data processing and covariates

Detailed data processing methods are in Appendix E. We regularised the two-dimensional positions to a 10-minute resolution using a continuous-time state-space model (Johnson *et al*., 2008; McClintock & Michelot, 2018), used linear interpolation to obtain an altitude for each location, and categorised each track as either a departure or return (Figure 3; Tarroux *et al*., 2016a). Each observed location was matched with 50 random points, with a gamma distribution of step lengths (parameters estimated from the empirical data) and uniform bearings. We used a dynamic elevation model (Liu *et al*., 2015) to account for land barriers by creating a categorical variable to denote whether each location was above or below ground (denoted *G*). Although no observed petrel locations were underground, random points were not spatially constrained, and this variable was designed to down-weight any impossible random points. We obtained hourly ERA5 wind vectors, composed of easting/northing wind at 14 altitudes (obtained from pressure). Note that the wind vectors are three-dimensional in the sense that they are altitude-dependent, but are not three-dimensional vectors themselves. We also obtained ground wind vectors (i.e., the vectors at the ground level from Liu *et al*., 2015). We also calculated wind support as the projection of the wind vector onto the vector of bird movement relative to airflow (see details in Tarroux *et al*., 2016a). This variable measures whether a bird is moving “with” the wind (positive support) or “against” the wind (negative support), and it is related to the energetic optimality of movement.

**Figure 3:**
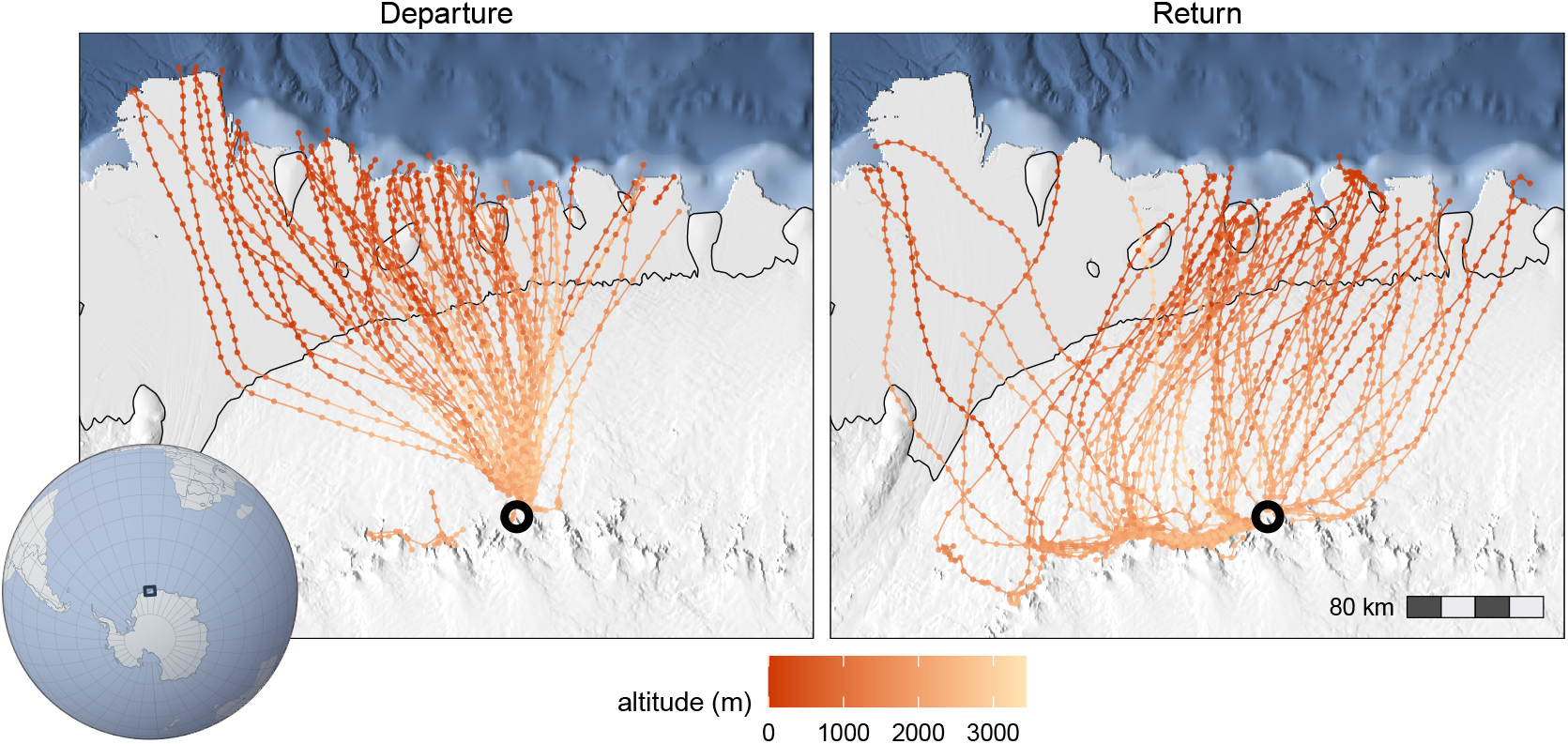
Map of petrel locations during departure and return trips, coloured by altitude. Open circle is the location of the petrel colony. Rectangle in globe represents study area, black line is the land border, and white beyond the land border is the Antarctic ice sheet.

### 5.2 Choice of movement kernel

Our first goal was to compare several potential three-dimensional movement kernels (described in Sections 3 and 4.1): i) a vMF (isotropic) CRW, ii) a Kent CRW, and iii) a BCRW with a bias towards small vertical bearings (i.e., level movement). In all CRWs, we modelled step lengths with a gamma distribution (by including the step length and its log), and included *G* to denote whether a step goes underground. The candidate models had the following SSF linear predictors:

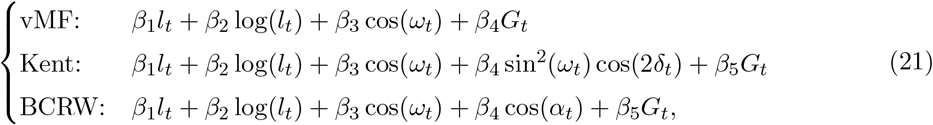

where the parameters for step length are related to the shape *a* and scale *b* of a gamma distribution through *β*_1_ = *a*− 3 and *β*_2_ = −1*/b*. For the vMF and Kent, *β*_3_ = *κ* is the concentration parameter. In the Kent SSF, ovalness is captured by *β*_4_ = *ρ*. In the BCRW, *β*_3_ and *β*_4_ measure the relative attractiveness of directional persistence and small vertical bearings, respectively. In all models, the coefficient for *G*_*t*_ measures selection for being underground; we did not intend to interpret this parameter (although we expect it to be very negative) but it was included to account for land’s barrier to movement.

We compared all models with AIC, which indicated that the top model was the BCRW (AIC = 3233), followed by the Kent (AIC = 8872) and isotropic (AIC = 9272) CRWs. We also generated tracks from each fitted movement kernel (using the simulation algorithm in Appendix D.1) to visually compare the empirical data to simulated data (Fieberg *et al*., 2024). This visual examination supported AIC results, where the geodesics simulated from the BCRW had a very similar distribution to the empirical data (with extreme ovaless; Figure 4). In the BCRW, we estimated a gamma distribution of step lengths with shape *â* = 4.47 and scale 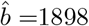 (i.e., mean 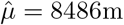, standard deviation 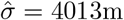). This example illustrates the flexibility of the BCRW formulation to capture the discrepancy in horizontal and vertical scales often found in three-dimensional movement data.

**Figure 4:**
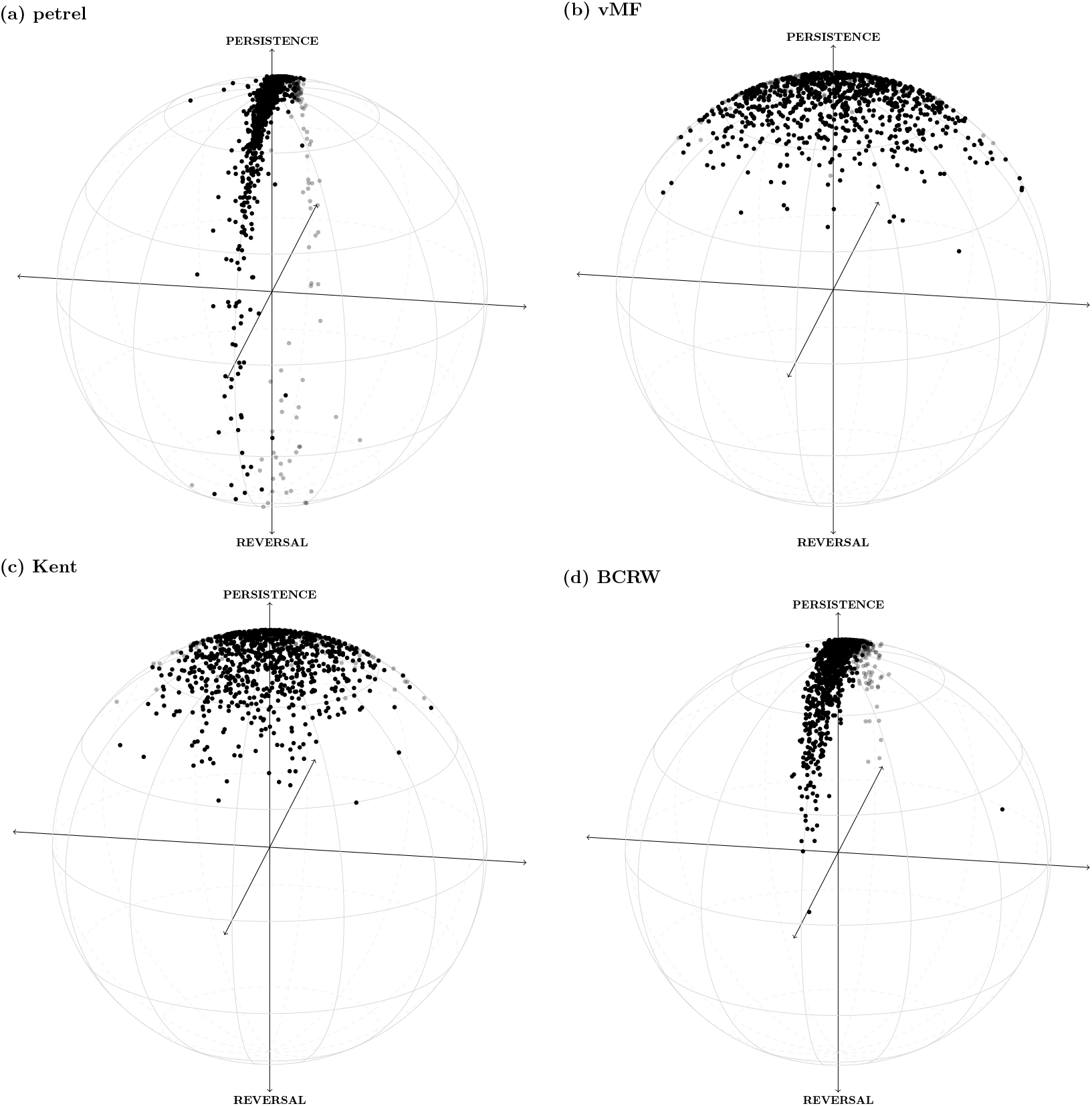
Plots of directional changes **u** on a unit sphere. (a) is the observed petrel data. (b-c) are simulated data from SSFs with the following movement kernels: (b) isotropic/vMF CRW; (c) Kent CRW; and (d) BCRW with selection for vertical bearing.

### 5.3 Ground vs. altitude-dependent wind

In the second analysis, we assessed how petrel movement decisions were influenced by external wind conditions. We expanded the BCRW of Section 5.2 to include wind support as a habitat variable. We interacted wind support with trip phase to assess how energetic strategies varied between departures and returns. To assess the effect of ignoring the vertical dimension of habitat, we compared models with wind support calculated from: i) ground wind vectors, or ii) altitude-dependent wind vectors. Therefore, the model (i) ignored vertical habitat structure while model (ii) integrated three-dimensional wind information.

AIC supported the model with altitude-dependent wind support (AIC = 3210) over the model with ground wind support (AIC = 3216). Both models indicated that petrels selected for positive wind support during departure trips, but the strength of the relationship was stronger for the ground wind model (Figure 5). Both models also had a negative interaction effect, where the selection for wind support was lower during return trips. However, the ground wind model indicated that petrels selected for negative wind support, suggesting that they tended to move against winds when returning to the colony. This was not the finding in the model with vertically-stratified wind, which indicated that there was more likely a small positive effect (Figure 5). Consistent with findings from Tarroux *et al*. (2016a), this suggests that petrels may select for energetically-optimal routes by varying their vertical position. This finding would not have been possible without considering the vertical structure of habitat, highlighting the ecological importance of modelling movement in three dimensions for flying species.

**Figure 5:**
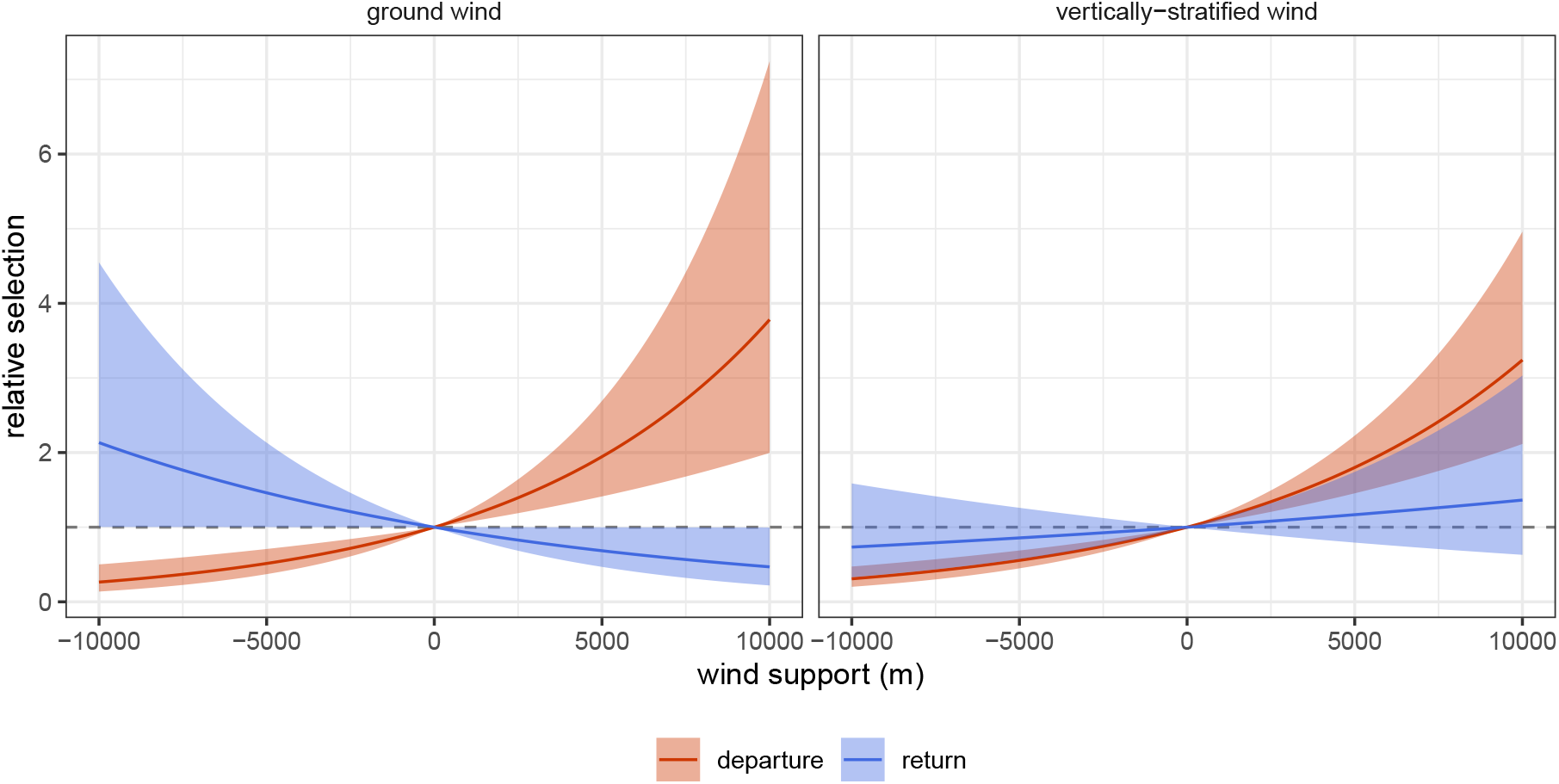
Relative selection strength for wind support (interacted with trip phase) in the second petrel analysis. Results are presented separately for the model based on ground wind (left) and the model based on vertically-stratified wind measured at the altitude of the birds (right). Confidence bands were obtained by generating posterior samples of all model parameters many times, generating predicted relation selection curves, and then taking pointwise 95% quantile intervals over a grid of wind support values.

## 6 Discussion

In this paper, we developed three-dimensional SSFs capable of jointly estimating movement and habitat selection, which can be easily fitted as conditional logistic regression using various software packages (e.g., Wood, 2017; Signer *et al*., 2019; Muff *et al*., 2020; Therneau, 2022). Our approach tackles unique challenges of three-dimensional data, via CRWs capable of capturing realistic patterns of displacement, directional changes, and angular biases. Therefore, our models improve upon: i) three-dimensional CRWs by increasing their flexibility and adding a spatially-explicit component; and ii) habitat selection models by porting the utility and flexibility of modern SSFs to a three-dimensional context.

Three-dimensional SSFs have great potential to better understand the spatial ecology of animals that swim, fly, burrow, or traverse complex terrain (Lennox *et al*., 2024). For these species, selection for three-dimensional variables is a better representation of how animals interact with their environment. In our example of Antarctic petrels, we showed how ignoring the third dimension of habitat can alter selection parameters and obscure behavioural and energetic patterns. We found that petrels may choose altitudes with more favourable wind conditions for energetic optimality during flight (consistent with Tarroux *et al*., 2016a); results which would have been missed with two-dimensional methods. Three-dimensional SSFs can also model how an animal’s vertical proximity to a variable affects their response to it (e.g., an animal at the ocean’s surface is unlikely to respond to features of the ocean floor; Lennox *et al*., 2024). The spatiotemporal nature of SSFs makes them a natural choice to understand between-animal interactions or other encounters (Potts & Börger, 2022), and two-dimensional methods may overestimate overlap if animals are vertically-stratified. For example, an animal at depth would not be at risk of collision with ships, even if they overlap in two-dimensional space and time. Therefore, three-dimensional SSFs may be important to understand niche partitioning (Lennox *et al*., 2024), overlap between individuals (Chandler *et al*., 2020), and encounter rates (Gurarie & Ovaskainen, 2013).

A realistic movement model is crucial for accurate estimation of habitat selection parameters (Forester *et al*., 2009), and subsequent spatial predictions (Potts & Börger, 2022). Therefore, we developed a very general family of three-dimensional CRWs aimed at capturing common features of three-dimensional data, such as anisotropic directional changes and biases. Isotropic models (i.e., the vMF CRW) are likely to allow for large vertical changes, which may be unrealistic in many studies, whereas the Kent CRW and BCRW have the capacity to constrain vertical movement. In the petrel example, extreme anisotropy was best captured by a BCRW with bias toward small vertical bearings. Although we focused on this particular anisotropic case where movement is mainly level, the Kent CRW or BCRW could accommodate other movement patterns. Future work could explore other spherical distributions (e.g., the extreme Kent distribution; Kent *et al*., 2018), non-parametric distributions (Klappstein *et al*., 2024; Arce Guillen *et al*., 2024), or position-based CRWs (Jonsen *et al*., 2005) for additional flexibility.

BCRWs have been hailed as some of the most flexible discrete-time movement models (Benhamou, 2014), and until now, they had seldom been explored in the three-dimensional context (but see BRWs in Ahmed *et al*., 2021). We focused on using BCRWs to capture anisotropy directional changes, but this approach can model bias relative to any spatial feature. In an SSF context, a BCRW provides a way to jointly model local habitat selection with larger-scale navigational cues (Benhamou, 2014; Duchesne *et al*., 2015). This could be important to capture navigation to and from a central place (e.g., foraging, haul-out, or nesting sites; McClintock *et al*., 2012; Tarroux *et al*., 2016a; Michelot *et al*., 2017), general home ranging behaviour (Signer *et al*., 2017), or between-patch movements (Duchesne *et al*., 2015). Our approach provides a general way to include height-dependent targets in SSFs, and this could be extended to include biases that are not simply towards or away from a target, to capture three-dimensional menotaxis in an SSF framework (Togunov *et al*., 2021).

Our proposed approach is an early step in three-dimensional movement modelling, and we are currently limited to systems where three-dimensional movement and environmental data has been or can be collected. Our model assumes that locations are regularly spaced in time, with little to no measurement error, and there are no existing tools to impute noisy or missing locations simultaneously for the horizontal and vertical dimensions. Future studies could focus on formulating three-dimensional CRWs as state-space models to account for measurement error (potentially uneven between dimensions), which could be used for data imputation (i.e., similar to Johnson *et al*., 2008, in two dimensions). We hope that this work motivates further collection of three-dimensional data, which which can be used to answer important ecological questions. Our approach directly builds on two-dimensional SSFs, and so it could be combined with many existing SSF extensions or implementations (e.g., non-linear and random effects, machine learning, Bayesian inference; Muff *et al*., 2020; Klappstein *et al*., 2024; Forrest *et al*., 2025). By more accurately representing how animals move in water or air, three-dimensional SSFs may lead to better inferences into and predictions of energetics (Eisaguirre *et al*., 2020; Klappstein *et al*., 2022), space use (Signer *et al*., 2017; Potts & Börger, 2022; Signer *et al*., 2024), behaviour (e.g., via three-dimensional HMM-SSFs Nicosia *et al*., 2017; Klappstein *et al*., 2023), and encounters (Noonan *et al*., 2021). It will important for future work to explore these methodological extensions, and their application to three-dimensional systems.

## Acknowledgements

We are grateful to Tarroux *et al*. (2016b) for making their petrel data publicly available on the Movebank Data Repository. We would also like to thank Louis-Paul Rivest, Mike Dowd, John Fieberg and Rob Lennox for discussions on three-dimensional modelling.

