## Appendices A-E for "Three-dimensional correlated random walks for animal movement and habitat selection"

### A Additional details for Section 2 (3D movement metrics)

#### A.1 Cartesian and polar/spherical coordinate systems

There are two coordinate systems used in this paper: Cartesian and spherical coordinates. In three dimensions, Cartesian coordinates are the signed distances of a point from each of perpendicular axes, whereas spherical coordinates describe that point in terms of a distance and two angles. Polar coordinates are the two-dimensional analogue of spherical coordinates. In movement ecology, we regularly toggle between these coordinate systems. For example, a two-dimensional animal location can be expressed as easting/northing (i.e., Cartesian) or latitude/longitude (spherical); this can be extended into three dimensions via the inclusion of altitude or depth measurements. Movement itself can be described as changes in easting, northing, and (potentially) height (i.e., Cartesian), or more commonly, in terms of a step length and bearings (i.e., spherical/polar) (Figure A1).

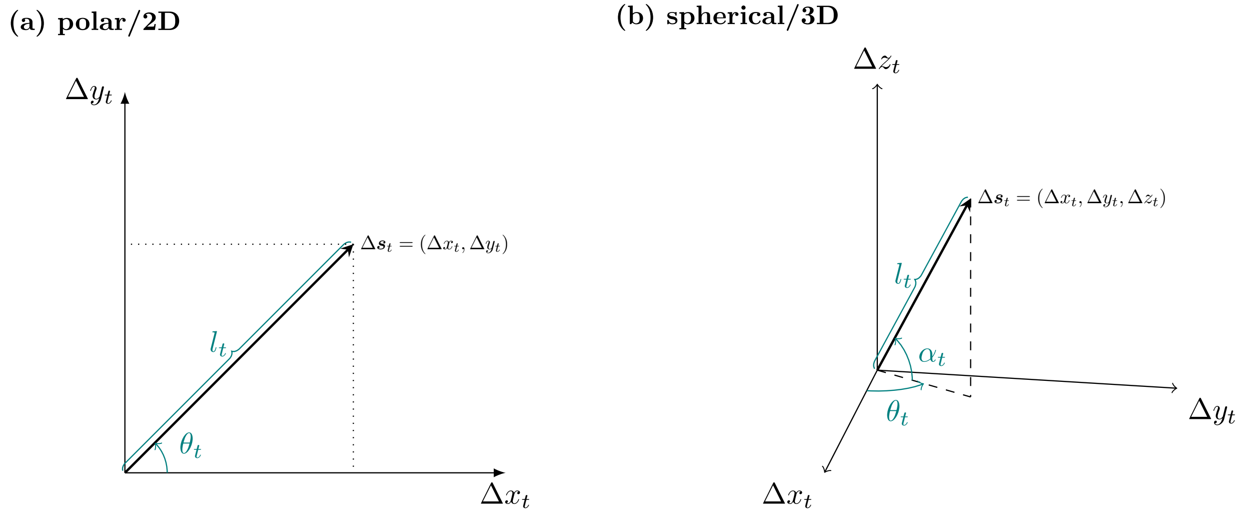

***Figure A1:*** *Illustration of polar and spherical coordinates of a movement step $\Delta \boldsymbol{s}_t$. In 2D (left), the step can be described by Cartesian coordinates $(\Delta x_t, \Delta y_t)$ or by polar coordinates $l_t$ (step length) and $\theta_t$ (bearing). In 3D (right), the Cartesian coordinates are $(\Delta x_t, \Delta y_t, \Delta z_t)$ and the spherical coordinates are $l_t$ (step length), $\theta_t$ (horizontal/azimuthal bearing) and $\alpha_t$ (vertical/polar bearing).*

##### A.1.1 Unit circle and unit sphere

The coordinates above can be scaled by their step length to obtain angular variables on a unit circle or sphere. This is a concept crucial to understanding the spherical modelling used in this paper. In this section, we provide details for these representations, first developing intuition for the two-dimensional case before expanding to the three-dimensional case used in our modelling. Note that we use bearings as an example to develop intuition, but the unit circle or sphere can also be used to plot directional changes.

In two dimensions, the direction of movement is measured by a single angular variable $\theta_t$. There are some challenges in working with angles; for example, $\theta_t = -\pi$ and $\theta_t = \pi$ correspond to the same direction. To address this circular nature, it is common to represent an angle on a unit circle (i.e., a circle of radius 1), where the angle is measured relative to East (Figure A2a). In this representation, it is clear that $\theta_t = -\pi$ and $\theta_t = \pi$ correspond to the same direction (West), because they are transformed to the same point on the unit circle: $\boldsymbol{v}_t = (-1, 0)$. A single angle $\theta_t$ can be transformed into two Cartesian coordinates $\boldsymbol{v}_t = (\cos(\theta_t), \sin(\theta_t))$, which are constrained such that $\boldsymbol{v}_t$ lies on the unit circle (Figure A2a).

In three dimensions, direction is measured by two angles $\theta_t$ and $\alpha_t$, and it can be represented as a point on the unit sphere (Figure A2b). Like in two dimensions, this is a convenient way to think about direction, because it clearly shows which directions are similar to each other (which isn't always obvious by looking at the values of $\alpha_t$ and $\theta_t$ directly). In this system, the direction $(\theta_t, \alpha_t)$ can then be equivalently described by a point on the surface of the sphere, with Cartesian coordinates,

$$\boldsymbol{v}_t = \begin{pmatrix} \cos(\alpha_t)\cos(\theta_t) \\ \cos(\alpha_t)\sin(\theta_t) \\ \sin(\alpha_t) \end{pmatrix}. \quad (1)$$

The unit circle or sphere can just as easily be used to show directional changes (i.e., turning angles or geodesics), but we use a different frame of reference for $\omega_t$. We show examples of this in the main-text (Figure 2).

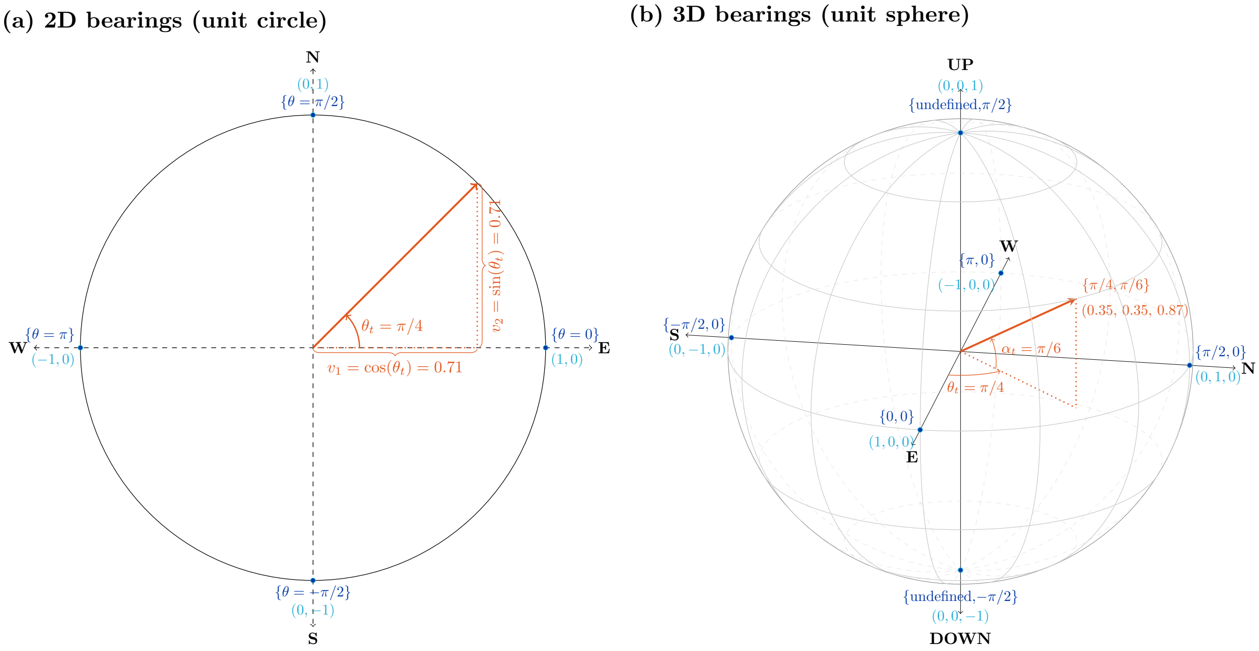

***Figure A2:*** *Example of to show a unit circle and unit sphere. (a) shows two-dimensional bearings on a unit circle, where the bearing $\theta_t$ is measured relative to East and the Cartesian coordinates are along the unit circle. (b) shows three-dimensional bearings on a unit sphere, where $\theta_t$ is measured relative to East and $\alpha_t$ is measured relative to the horizontal plane. The Cartesian coordinate is on the surface of the unit sphere, measured as the distances from each axis.*

#### A.2 Justification for the geodesic

Consider two successive movement directions $(\theta_{t-1}, \alpha_{t-1})$ and $(\theta_t, \alpha_t)$. A naive approach to calculate the change in direction is to take differences for the two angles: $\theta_t^* = \theta_t - \theta_{t-1}$ and $\alpha_t^* = \alpha_t - \alpha_{t-1}$ (i.e., "turning angles"). Alternatively, the approach we advocate for is to translate the angles into two directions $\boldsymbol{v}_{t-1}$ and $\boldsymbol{v}_t$ on the unit sphere, and measure the directional change as the shortest arc connecting $\boldsymbol{v}_{t-1}$ to $\boldsymbol{v}_t$ (i.e., the geodesic). Here, we use an example to compare the two methods, and explain why the naive approach fails to capture the spherical nature of the process.

Figure A3 shows two example scenarios. In one of them (shown in teal), an animal is moving completely horizontally, and goes from moving due North at time 1 to moving due South at time 2. This corresponds to a complete reversal in direction, as can be seen in the figure where $\boldsymbol{v}_1$ and $\boldsymbol{v}_2$ are on opposite sides of the sphere. In the second scenario (shown in purple), an animal is moving almost only vertically (upwards), but changes from moving slightly towards the North at time 1 to moving slightly towards the South at time 2. This is a very small change in direction, and, in this scenario, $\boldsymbol{v}_1$ and $\boldsymbol{v}_2$ are very close to each other on the sphere. The naive approach yields the same results for the two scenarios because, in each case, the animal's vertical bearing doesn't change ($\alpha_t^* = \alpha_t - \alpha_{t-1} = 0$), and the horizontal bearing changes completely from North to South ($\theta_t^* = \theta_t - \theta_{t-1} = \pi$). We see that this is not an appropriate representation of the directional change. On the other hand, the geodesic approach is much more informative, as it correctly captures the fact that the first scenario has a much larger change in direction (arc length $\omega = \pi$) than in the second scenario ($\omega = 0.16$).

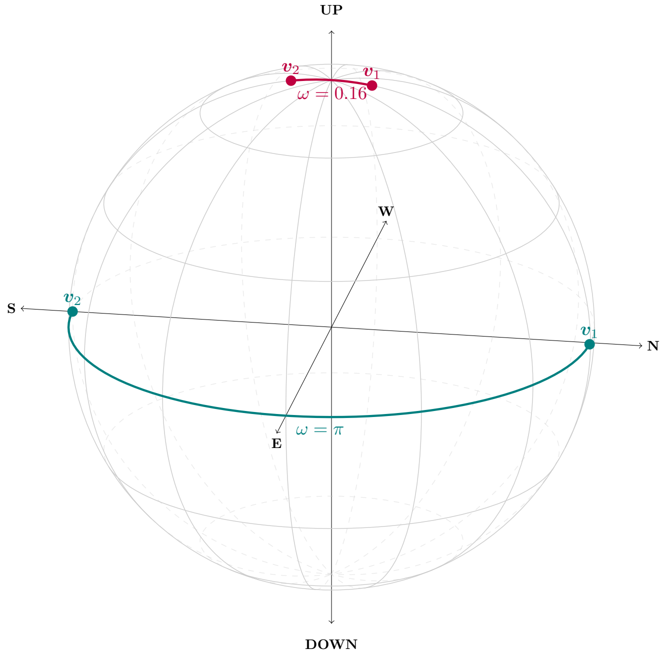

***Figure A3:*** *Example to illustrate the utility of geodesics to represent changes in direction. Two scenarios are shown: one where the animal's direction reverses completely (teal), and one where the direction barely changes (purple). First-order differences in horizontal and vertical bearings are identical for the two scenarios, whereas the geodesics (coloured lines) capture the difference between the two scenarios.*

#### A.3 Tables of important transformations

***Table 1:*** *Summary of spherical coordinates used in the main text. Note that* $\Delta$ *refers to the difference between the previous and current Cartesian coordinate, e.g.,* $\Delta x_t = x_t - x_{t-1}$.

| **Spherical coordinate** | **Definition** | **Range** |
|---|---|---|
| step length | $l_t = \sqrt(\Delta x_t^2 + \Delta y_t^2 + \Delta z_t^2)$ | $l_t \in [0, \infty)$ |
| horizontal bearing | $\theta_t = \text{atan}_2[\Delta y_t, \Delta x_t]$ | $\theta_t \in [-\pi, \pi)$ |
| vertical bearing | $\alpha_t = \text{atan}_2[\sqrt{\Delta x_t^2 + \Delta y_t^2}, \Delta z_t]$ | $\alpha_t \in [-\pi/2, \pi/2]$ |
| horizontal turning angle | $\theta_t^* = \theta_t - \theta_{t-1}$ | $\theta_t^* \in [-\pi, \pi)$ |
| vertical turning angle | $\alpha_t^* = \alpha_t - \alpha_{t-1}$ | $\alpha_t^* \in [-\pi, \pi)$ |
| arc size | $\omega_t = \cos^{-1}[\sin(\alpha_{t-1})\sin(\alpha_t) + \cos(\alpha_{t-1})\cos(\alpha_t)\cos(\theta_t - \theta_{t-1})]$ | $\omega_t \in [0, \pi]$ |
| initial orientation | $\delta_t = \text{atan}_2[\sin(\alpha_t) - \cos(\omega_{t-1})\sin(\alpha_{t-1}),\ \cos(\alpha_{t-1})\cos(\alpha_t)\sin(\theta_t - \theta_{t-1})]$ | $\delta_t \in [-\pi, \pi)$ |

***Table 2:*** *Summary of Cartesian coordinates used in this chapter, and their relationship to spherical coordinates* $L_t, \theta_t, \alpha_t\omega_t, \delta_t$. *Note that the trigonometric functions are swapped for the vertical bearing* $\alpha_t$ *(compared to standard spherical-Cartesian transformations) because it is measured relative to horizontal rather than relative to "up".*

| **Cartesian coordinate** | **Definition (from** $L_t, \theta_t, \alpha_t, \omega_t, \delta_t$**)** | **Range** |
|---|---|---|
| geographic position | $\boldsymbol{s}_t = \begin{pmatrix} x_t \\ y_t \\ z_t \end{pmatrix} = \begin{pmatrix} x_{t-1} + l_t \sin(\alpha_t)\cos(\theta_t) \\ y_{t-1} + l_t \sin(\alpha_t)\sin(\theta_t) \\ z_{t-1} + l_t \cos(\alpha_t) \end{pmatrix}$ | $\boldsymbol{s}_t \in \mathbb{R}^3$ |
| bearings | $\boldsymbol{v}_t = \begin{pmatrix} v_{1,t} \\ v_{2,t} \\ v_{2,t} \end{pmatrix} = \begin{pmatrix} \cos(\alpha_t)\cos(\theta_t) \\ \cos(\alpha_t)\sin(\theta_t) \\ \sin(\alpha_t) \end{pmatrix}$ | $\boldsymbol{v}_t \in [-1, 1]^3$ |
| turning angles | $\boldsymbol{u}_t = \begin{pmatrix} u_{1,t} \\ u_{2,t} \\ u_{2,t} \end{pmatrix} = \begin{pmatrix} \sin(\omega_t)\cos(\delta_t) \\ \sin(\omega_t)\sin(\delta_t) \\ \cos(\omega_t) \end{pmatrix}$ | $\boldsymbol{u}_t \in [-1, 1]^3$ |

### B Additional details for Section 3 (3D CRWs)

#### B.1 vMF and Kent additional details

The von Mises Fisher (vMF) distribution is a spherical distribution, characterised by two parameters: the mean direction $\boldsymbol{\mu}$ and concentration $\kappa$. $\kappa$ describes the variance around the mean. In our case, we are modelling the unit vector $\boldsymbol{u}$ of geodesic coordinates, and we usually assume that animals exhibit directional persistence $\boldsymbol{\mu} = (0, 0, 1)$ (Figure B1a). If $\kappa$ is not constrained to be positive, a negative value corresponds to a mean direction of $\boldsymbol{\mu} = (0, 0, -1)$, corresponding to reversals in direction.

The Kent distribution is a generalisation of the vMF, where we allow anisotropy via the specification of an ellipse. The orientation of this ellipse is controlled by the mean direction $\boldsymbol{\mu}$, a major axis that runs along the length of the ellipse, and a minor axis that is perpendicular to the major. The distribution is also controlled by a concentration parameter $\kappa$ and an ovalness parameter $\rho$. In our case, we are most likely to assume that a major axis that can capture a tendency to make larger horizontal directional changes, compared to vertical (Figure B1b). If the ovalness parameter $\rho$ is negative, this swaps the major and minor axes of the distributions, which can be useful in systems where changes in direction are predominantly vertical.

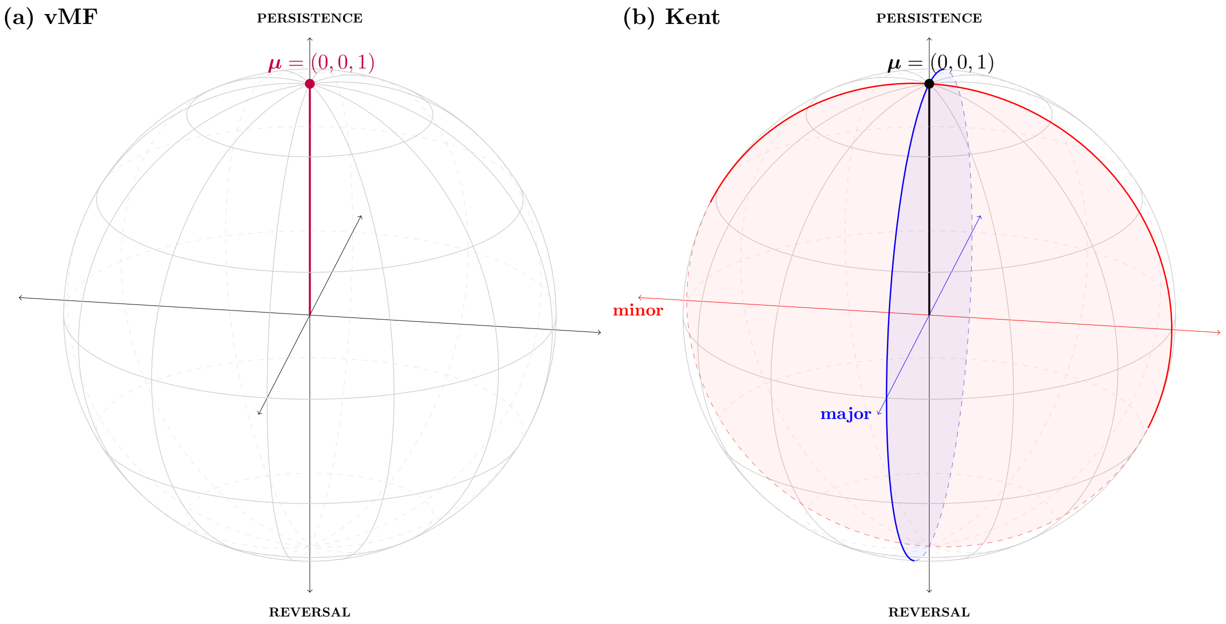

***Figure B1:*** *Example of von Mises Fisher (vMF) and Kent distribution parameters describing orientation, under our chosen frame of reference. (a) vMF with mean direction* $\boldsymbol{\mu}$ *set to (0, 0,1) to capture directional persistence. (b) Kent with* $\boldsymbol{\mu}$ *set to (0, 0,1) to capture directional persistence, and major/minor axes.*

#### B.2 BCRW additional details

Here, we present how to extend the CRW to include any number of directional biases. Consider we want to model an animal's direction at time $t$ as a compromise between $K$ directional targets, one of which is the previous direction. Following Duchesne et al. (2015), we can write these directional targets as unit vectors, $\widetilde{\gamma}_{1t}, \widetilde{\gamma}_{2t}, \dots \widetilde{\gamma}_{Kt}$, and the overall mean direction is $\boldsymbol{\gamma}_t = \kappa_1 \widetilde{\gamma}_{1t} + \kappa_2 \widetilde{\gamma}_{2t} + \dots + \kappa_K \widetilde{\gamma}_{Kt}$. We want to model the direction of travel as a vMF distribution, based on the "consensus" model approach, such that

$$f(\boldsymbol{v}_t) = C_p(\kappa_t) \exp(\kappa_t \boldsymbol{\mu}_t^\top \boldsymbol{v}_t) \tag{2}$$

where the concentration parameter is $\kappa_t = ||\boldsymbol{\gamma}_t||$ and the mean direction $\boldsymbol{\mu}_t$ is obtained by scaling $\boldsymbol{\gamma}_t$ to a unit vector, $\boldsymbol{\mu}_t = \frac{\boldsymbol{\gamma_t}}{||\boldsymbol{\gamma}_t||}$. Plugging this into Equation 2 (and ignoring the normalising constant for now), we get

$$\begin{align}
f(\boldsymbol{v}_t) &\propto \exp(||\boldsymbol{\gamma}_t|| \frac{\boldsymbol{\gamma_t}}{||\boldsymbol{\gamma}_t||}^\top \boldsymbol{v}_t) \tag{3}\\
&\propto \exp(\boldsymbol{\gamma_t}^\top \boldsymbol{v}_t) \tag{4}\\
&\propto \exp([\kappa_1 \widetilde{\gamma}_{1t} + \kappa_2 \widetilde{\gamma}_{2t} + \dots + \kappa_K \widetilde{\gamma}_{Kt}]^\top \boldsymbol{v}_t) \tag{5}\\
&\propto \exp(\kappa_1 \widetilde{\gamma}_{1t}^\top \boldsymbol{v_t} + \kappa_2 \widetilde{\gamma}_{2t}^\top \boldsymbol{v_t} + \dots \kappa_K \widetilde{\gamma}_{Kt}^\top \boldsymbol{v_t}) \tag{6}\\
&\propto \exp(\kappa_1 \cos(\omega_{1t}) + \kappa_2 \cos(\omega_{2t}) + \dots + \kappa_K \cos(\omega_{Kt})), \tag{7}
\end{align}$$

where $\omega_{it}$ is the arc lengths between $\boldsymbol{v}_t$ and $\gamma_{it}$. The simplification in the last step comes from the inner product rule.

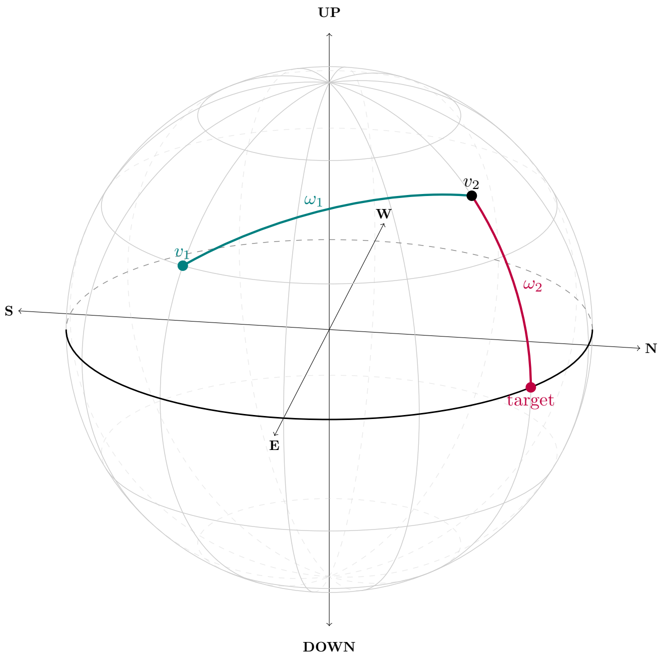

***Figure B2:*** *Example of a BCRW with bias towards level bearings ($\omega_1$ is the arc length between successive bearings, and $\omega_2$ is the arc length between the second bearing and a target of $\alpha = 0$).*

### C Additional details for Section 4 (3D SSF)

#### C.1 Change of variable

The change of variable from $f(L, \omega, \delta)$ to $f(x, y, z)$ is,

$$f(x,y,z) = \frac{1}{|\det(\mathbf{J})|} f(L,\omega,\delta) \tag{8}$$

where $\mathbf{J}$ is the Jacobian matrix of the transformation between spherical and Cartesian coordinates. First, we can define the relationship between $(x, y, z)$ and $(L, \omega, \delta)$ via the following functions,

$$x = L\sin(\omega)\cos(\delta) = v_1(L,\omega,\delta) \tag{9}$$

$$y = L\sin(\omega)\sin(\delta) = v_2(L,\omega,\delta) \tag{10}$$

$$z = L\cos(\omega) = v_3(L,\omega). \tag{11}$$

This has the corresponding $3 \times 3$ Jacobian matrix,

$$\mathbf{J} = \begin{bmatrix} \frac{\partial v_1}{\partial L}(L,\omega,\delta) & \frac{\partial v_1}{\partial \omega}(L,\omega,\delta) & \frac{\partial v_1}{\partial \delta}(L,\omega,\delta) \\ \frac{\partial v_2}{\partial L}(L,\omega,\delta) & \frac{\partial v_2}{\partial \omega}(L,\omega,\delta) & \frac{\partial v_2}{\partial \delta}(L,\omega,\delta) \\ \frac{\partial v_3}{\partial L}(L,\omega) & \frac{\partial v_3}{\partial \omega}(L,\omega) & \frac{\partial v_3}{\partial \delta}(L,\omega) \end{bmatrix} \tag{12}$$

$$= \begin{bmatrix} \sin(\omega)\cos(\delta) & L\cos(\omega)\cos(\delta) & -L\sin(\omega)\sin(\delta) \\ \sin(\omega)\sin(\delta) & L\cos(\omega)\sin(\delta) & L\sin(\omega)\cos(\delta) \\ \cos(\omega) & -L\sin(\omega) & 0 \end{bmatrix} \tag{13}$$

with the corresponding determinant $|\det(\mathbf{J})| = L^2 \sin(\omega)$ for $0 < \omega \leq \pi$ . Therefore, the three-dimensional spatial distribution implied from the distributions of step lengths and turning angles is,

$$f(x,y,z) = \frac{1}{L^2\sin(\omega)} f(L,\omega,\delta). \tag{14}$$

The $L^2$ term in the denominator can be viewed as a correction for the fact that the volume of a spherical shell of radius $L$ increases as a function of $L^2$. The $\sin(\omega)$ term accounts for the fact that the circumference of a circle of constant $\omega$ on the unit sphere is proportional to $\sin(\omega)$.

#### C.2 Random sampling

##### C.2.1 Uniform Monte Carlo

A naive choice of $h$ would be to sample spatially uniform points, such that $h(\boldsymbol{s}_t, \boldsymbol{r}_{it})$ is constant for all points (i.e., uniform Monte Carlo; Michelot et al., 2024). This has been previously discussed for two-dimensional SSFs, where it is suggested to sample uniform points on a disc (centred on the previous location) large enough to encompass all substantial probability density (Figure C1; Klappstein et al., 2022; Michelot et al., 2024). In three dimensions, this can be extended to sampling uniformly within a sphere centred on $\boldsymbol{s}_t$ with radius $R$. To do so, uniform bearings can be generated by sampling points from a three-dimensional Gaussian distribution (i.e., each component is from $N(0, 1)$), which are then scaled by their length to obtain points on the unit sphere $\boldsymbol{v}_{it}$. Step lengths are not drawn from a uniform distribution directly, as this would lead to more points near the centre of the sphere. Instead, we can sample each step length as the cubic root of a random draw from $\text{Unif}(0, R^3)$.

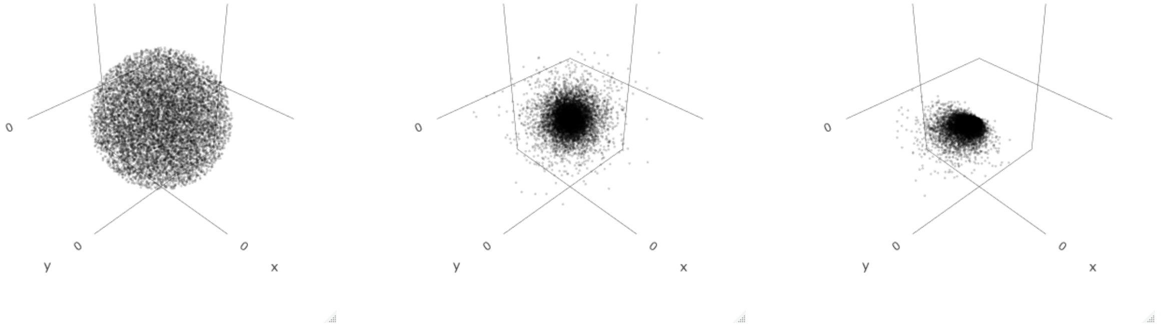

***Figure C1:*** *Examples of random/integration points, centred on point (0, 0, 0). Left panel is spatially uniform integration points within a sphere (i.e., uniform Monte Carlo). Middle panel shows points with gamma-distributed step lengths and uniform bearings. Right panel shows points with gamma-distributed step lengths and Kent-distributed arcs.*

##### C.2.2 Importance sampling

A more efficient option is to choose $h$ to be something that samples where the probability is expected to be highest (Michelot et al., 2024). We can leverage our knowledge of the animal's movement (i.e., its speed and directional persistence) to choose a sensible distribution. $h$ could take many forms, but it is convenient to use the same distributional form as $\phi$ (for implementation, described in the next section). Therefore, we can generate random points with step lengths that follow a gamma distribution, $\tilde{l} \sim \text{Gamma}(\tilde{a}, \tilde{b})$ where $\tilde{a}$ and $\tilde{b}$ are the shape and scale parameters, respectively. In this case, $h$ is the three-dimensional spatial distribution implied from a gamma distribution of step lengths. We could further improve the importance sampling by generating arcs (i.e., directional changes) from a Kent distribution (Figure C1).

##### C.3 Parameter corrections

If the SSF parameters are estimated via conditional logistic regression (CLR), then the estimated CLR parameters (denoted $\tilde{\boldsymbol{\beta}}$) will represent deviations from $h$, rather than the coefficients of interest (denoted $\boldsymbol{\beta}$). We can still fit the SSF as CLR, but we must update the estimated parameters to reflect our choice of $h$. Following Forester et al. (2009); Avgar et al. (2016), the appropriate corrections can be found by simplifying,

$$\frac{\phi(\boldsymbol{s}_t \mid \boldsymbol{s}_{1:(t-1)})}{h(\boldsymbol{s}_t \mid \boldsymbol{s}_{1:(t-1)})} = \frac{\exp\{\boldsymbol{\beta}^\top \boldsymbol{c}(\boldsymbol{s}_t \mid \boldsymbol{s}_{1:(t-1)})\}}{h(\boldsymbol{s}_t \mid \boldsymbol{s}_{1:(t-1)})} \tag{15}$$

where $h$ has parameters $\tilde{\boldsymbol{\theta}}$. Ultimately, we want to obtain $\boldsymbol{\beta}$ which can then be transformed to the parameters of the distributions $\boldsymbol{\theta}$.

**Gamma distribution:** Assume we sample random points with step lengths that follow a gamma distribution (i.e., $\tilde{l} \sim \text{Gamma}(\tilde{a}, \tilde{b})$). The corrections for to go from $\tilde{\boldsymbol{\beta}}$ to $\boldsymbol{\beta}$ are obtained as follows,

$$\frac{\phi(\boldsymbol{s}_t \mid \boldsymbol{s}_{1:(t-1)})}{h(\boldsymbol{s}_t \mid \boldsymbol{s}_{1:(t-1)})} = \frac{\exp\{\beta_1 l_t + \beta_2 \log(l_t)\}}{\exp\{(-1/\tilde{b}) l_t + (\tilde{a} - 3)\log(l_t)\}} \tag{16}$$

$$= \exp\{\beta_1 l_t + \beta_2 \log(l_t) - (-1/\tilde{b}) l_t - (\tilde{a} - 3)\log(l_t)\} \tag{17}$$

$$= \exp\{(\beta_1 + 1/\tilde{b}) l_t + (\beta_2 - \tilde{a} + 3)\log(l_t)\}. \tag{18}$$

Therefore,

$$\tilde{\beta}_1 = \beta_1 + 1/\tilde{b} \quad \Rightarrow \quad \beta_1 = \tilde{\beta}_1 - 1/\tilde{b} \tag{19}$$

$$\tilde{\beta}_2 = \beta_2 - \tilde{a} + 3 \quad \Rightarrow \quad \beta_2 = \tilde{\beta}_2 + \tilde{a} - 3. \tag{20}$$

Given the relationships $\beta_1 = -1/b$ and $\beta_2 = a - 3$, we can also obtain the step length parameters,

$$b = -\frac{1}{\tilde{\beta}_1 - 1/\tilde{b}}\ , \quad a = \tilde{\beta}_2 + \tilde{a}. \tag{21}$$

**Kent distribution:** The corrections for a Kent distribution can be obtained as follows,

$$\frac{\phi(\boldsymbol{s}_{t+1} \mid \boldsymbol{s}_{1:t})}{h(\boldsymbol{s}_{t+1} \mid \boldsymbol{s}_{1:t})} = \frac{\exp\{\beta_1 \cos(\omega_t) + \beta_2 \sin^2(\omega_t)\cos(2\delta_t)\}}{\exp\{\tilde{\kappa}\cos(\omega_t) + \tilde{\rho}\sin^2(\omega_t)\cos(2\delta_t)\}} \tag{22}$$

$$= \exp\{(\beta_1 - \tilde{\kappa})\cos(\omega_t) + (\beta_2 - \tilde{\rho})\sin^2(\omega_t)\cos(2\delta_t)\} \tag{23}$$

where $\tilde{\kappa}$ and $\tilde{\rho}$ are the parameters used to generate random arcs. Therefore, we know the relationship between $\tilde{\boldsymbol{\beta}}$ and $\boldsymbol{\beta}$ to be,

$$\tilde{\beta}_1 = \beta_1 - \tilde{\kappa} \quad \Rightarrow \quad \beta_1 = \tilde{\beta}_1 + \tilde{\kappa} \tag{24}$$

$$\tilde{\beta}_2 = \beta_2 - \tilde{\rho} \quad \Rightarrow \quad \beta_2 = \tilde{\beta}_2 + \tilde{\rho}. \tag{25}$$

Note the relationships $\beta_1 = \kappa$ and $\beta_2 = \rho$ to obtain the estimated Kent parameters. Also, importantly, the vMF is a special case of the Kent where $\rho = 0$.

### D Simulations

#### D.1 Simulation algorithm

All simulations use the same algorithm to simulate a track of length $T$ from an SSF with known parameters $\boldsymbol{\beta}$. First, generate random a random first step ending at $\boldsymbol{s}_0$ with a random step length and bearings. This won't be included in the final track but is needed to calculate directional changes. Then, at each iteration $i$, do the following,

1. For some large number $K$, generate proposed endpoints $\boldsymbol{r}_{i,k}$, $k \in \{1, 2, \ldots, K\}$ centred on $\boldsymbol{s}_{i-1}$. These could be generated uniformly within a sphere, but a more efficient method is to generate random points with importance sampling (i.e., from some distribution $g$). The most obvious choice for simulation is to generate points from the known movement kernel, where possible. For example, we can sample step lengths as $l_{i,k} \sim \text{Gamma}(a, b)$ with shape $a = \beta_{\log(l)} + 3$ and scale $b = -1/\beta_l$. Bearings can be generated in two ways:

   (a) Uniformly on a unit sphere, by simulating points from a three-dimensional Gaussian distribution. Assuming independence, each component is from $N(0, 1)$. This can be scaled by its length to obtain points on the unit sphere $\boldsymbol{v}_{i,k}$.

   (b) Based on geodesics from a Kent distribution. Geodesics are generated $\boldsymbol{u}_{i,k} \sim \text{Kent}(\kappa, \rho)$ and converted to bearings $\boldsymbol{v}_{i,k}$ following formulas in Benhamou (2019).

   Then, $g$ is the three-dimensional distribution of the step lengths and bearings. If spatially uniform, then $g$ is constant across all points.

2. Transform the step lengths and bearings to geographic positions: $\boldsymbol{s}_{i,k} = \boldsymbol{s}_{i-1} + L_{i,k}\boldsymbol{v}_{i,k}$.

3. Sample $\boldsymbol{s}_i$ from $\boldsymbol{r}$ with probabilities proportional to $p_{i,k}$, calculated based on the SSF,

$$p_{i,k} = \frac{\exp\{\boldsymbol{\beta}^\top \boldsymbol{c}(\boldsymbol{s}_{i-1}, \boldsymbol{r}_{i,k})\}/g(\boldsymbol{s}_{i-1}, \boldsymbol{r}_{i,k})}{\sum_{j=1}^{K} \exp\{\boldsymbol{\beta}^\top \boldsymbol{c}(\boldsymbol{s}_{i-1}, \boldsymbol{r}_{j,k})\}/g(\boldsymbol{s}_{i-1}, \boldsymbol{r}_{j,k})}. \tag{26}$$

   Note if the proposal points are sampled from the true movement kernel (i.e., $g = \phi$), the associated movement covariates cancel out from the linear predictor at this step (as explained in Forester et al., 2009).

If there are no habitat covariates, this algorithm reduces to simulating a three-dimensional CRW; this can be simulated with only one proposed endpoint per iteration if the endpoints are sampled from the true movement kernel.

#### D.2 Simulation to check SSF implementation

We conducted simulations to assess the accuracy and precision of our proposed SSF implementation, when fitting a correctly-specified model. The general simulation procedure was to: i) generate a movement track from an SSF with known parameters, ii) generate random/integration points from chosen distributions, and iii) fit a correctly-specified SSF to assess the variance and bias of the parameter estimators. We assessed SSF estimators for all three CRWs (isotropic, Kent, BCRW), presented in the main-text. Given that we expect that integration methods may be particularly important in higher dimensions, we contrasted the performance of uniform Monte Carlo and two importance sampling designs (described below).

**Methods** For each scenario, we conducted 100 simulation iterations, where in each we generated a track of length $T = 1000$ from an SSF with known parameters. All SSFs had the same step length distribution and habitat selection (described in point 1, below), but varied in their angular distributions. For each track, we generated three sets of random points: i) uniformly within a sphere with a radius equal to the maximum observed step length, ii) using importance sampling with gamma-distributed step lengths, and iii) same as (ii) but also with Kent-distributed arcs. We also tested performance with 25, 50, 100, 200, 500, and 1000 integration points for each method. We fitted each SSF as conditional logistic regression using the `survival` package and calculated relative error ($\hat{\beta} - \beta/\beta$) for each coefficient. We considered three main simulation scenarios:

1. **Isotropic CRW:** The simplest scenario was an isotropic SSF, with the following linear predictor:
$$\eta = \beta_1 L + \beta_2 \log(L) + \beta_3 \cos(\omega) + \beta_4 x_1 \tag{27}$$
where $\beta_1 = -0.5$, $\beta_2 = -1$, $\beta_3 = \kappa = 6$, and $\beta_4 = 3$. The step length coefficients correspond to a gamma-distribution with shape $a = 2$ and scale $b = 2$. The habitat covariate $x_1$ was created by generating uniform random noise over a 3D spatial grid, and then applying a moving average over a spherical window to obtain spatial autocorrelation, using the imager R package.

2. **Kent CRW:** We also assessed SSF estimators of the Kent CRW, with the linear predictor:
$$\eta = \beta_1 l + \beta_2 \log(l) + \beta_3 \cos(\omega) + \beta_4 x_1 + \beta_5 \sin^2(\omega)\cos(2\delta) \tag{28}$$
where $\beta_1, \beta_2, \beta_3, \beta_4$ are the same as the isotropic case, and $\beta_5 = \rho = 3$.

3. **BCRW:** Lastly, we assessed the SSF estimators of a BCRW with attraction towards level movement:
$$\eta = \beta_1 l + \beta_2 \log(l) + \beta_3 \cos(\omega) + \beta_4 x_1 + \beta_5 \cos(\theta) \tag{29}$$
where $\beta_1, \beta_2, \beta_3, \beta_4$ are the same as the isotropic/Kent scenarios, and $\beta_5 = \kappa_2 = 100$. Note that here $\beta_3$ is equal to $\kappa_1$. Further, in this case, we only compared two integration schemes: i) uniform Monte Carlo, and ii) importance sampling with gamma step lengths and uniform bearings. We did not assess the Kent distribution because we did not derive parameter corrections for Kent/vMF integration points and a BCRW.

**Results** In all scenarios, there was more variability in coefficients estimated with uniform Monte Carlo, particularly for step length parameters (Figures D1 - D3). There was no discernible bias for any of the coefficients, but there was generally greater variability in the habitat parameter. There was very little difference between the two importance sampling methods (Figures D1, D2), suggesting that sampling integration points with a step length distribution has a greater impact on parameter estimation than an angular distribution.

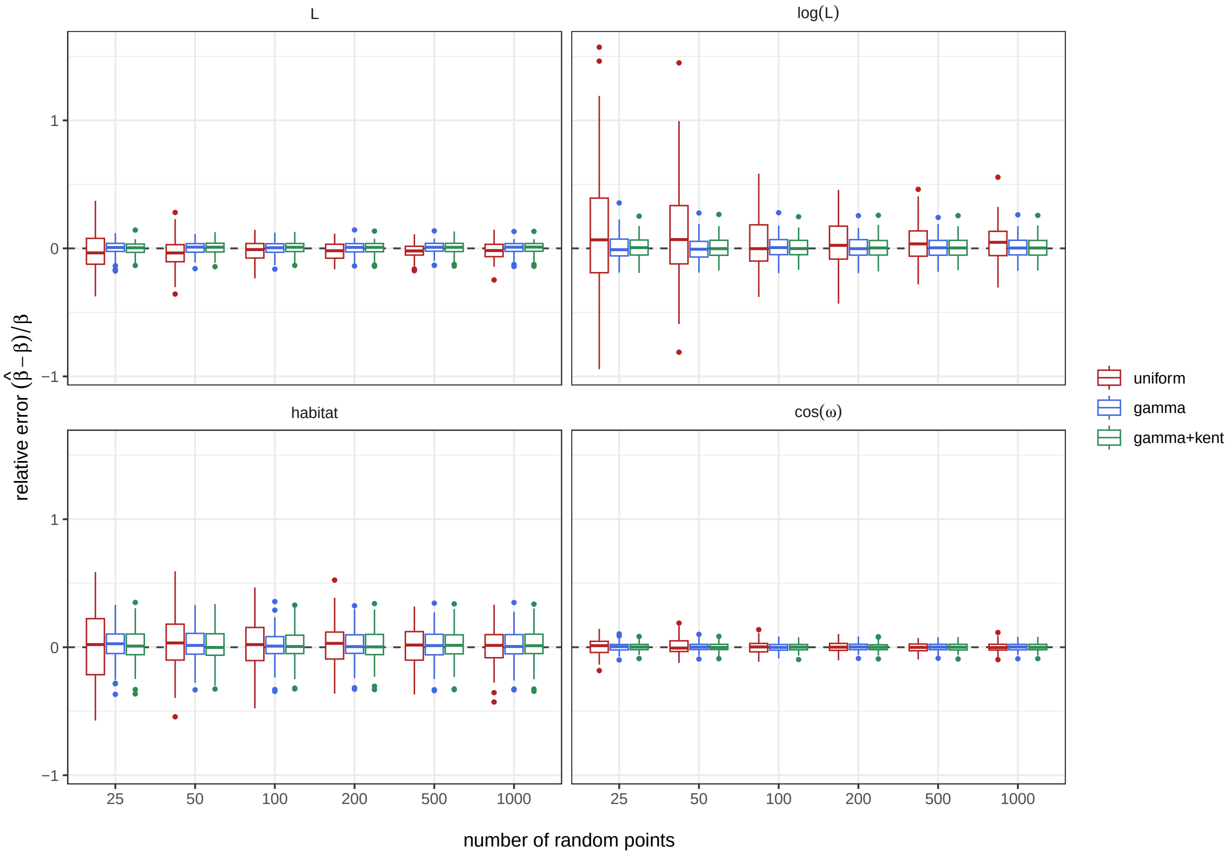

***Figure D1:*** *Results of the simulation study to check SSF implementation for the isotropic CRW. Note that the coefficient for* $\cos(\omega)$ *corresponds to* $\kappa$*.*

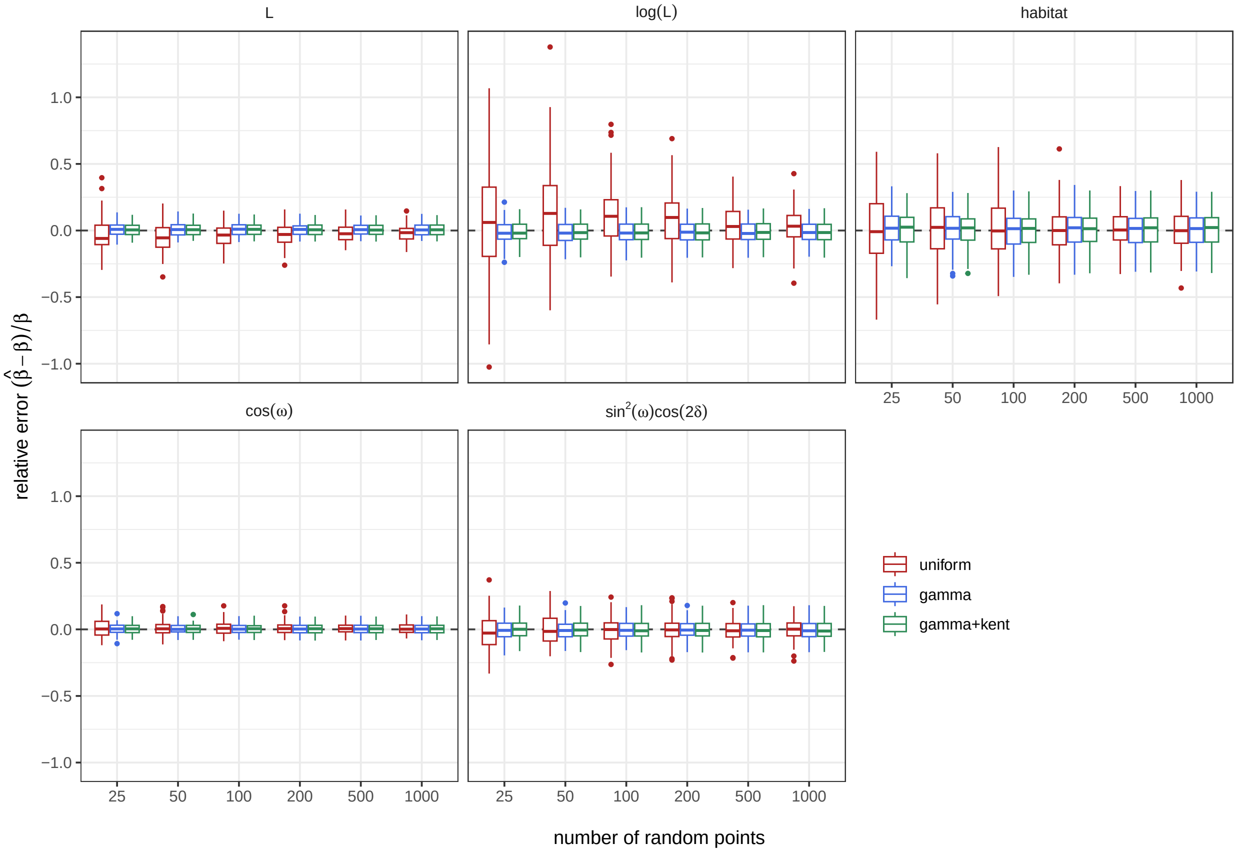

***Figure D2:*** *Results of the simulation study to check SSF implementation for the Kent CRW. Note that the coefficients for* $\cos(\omega)$ *and* $\sin^2(\omega)\cos(2\delta)$ *correspond to* $\kappa$ *and* $\rho$ *directly.*

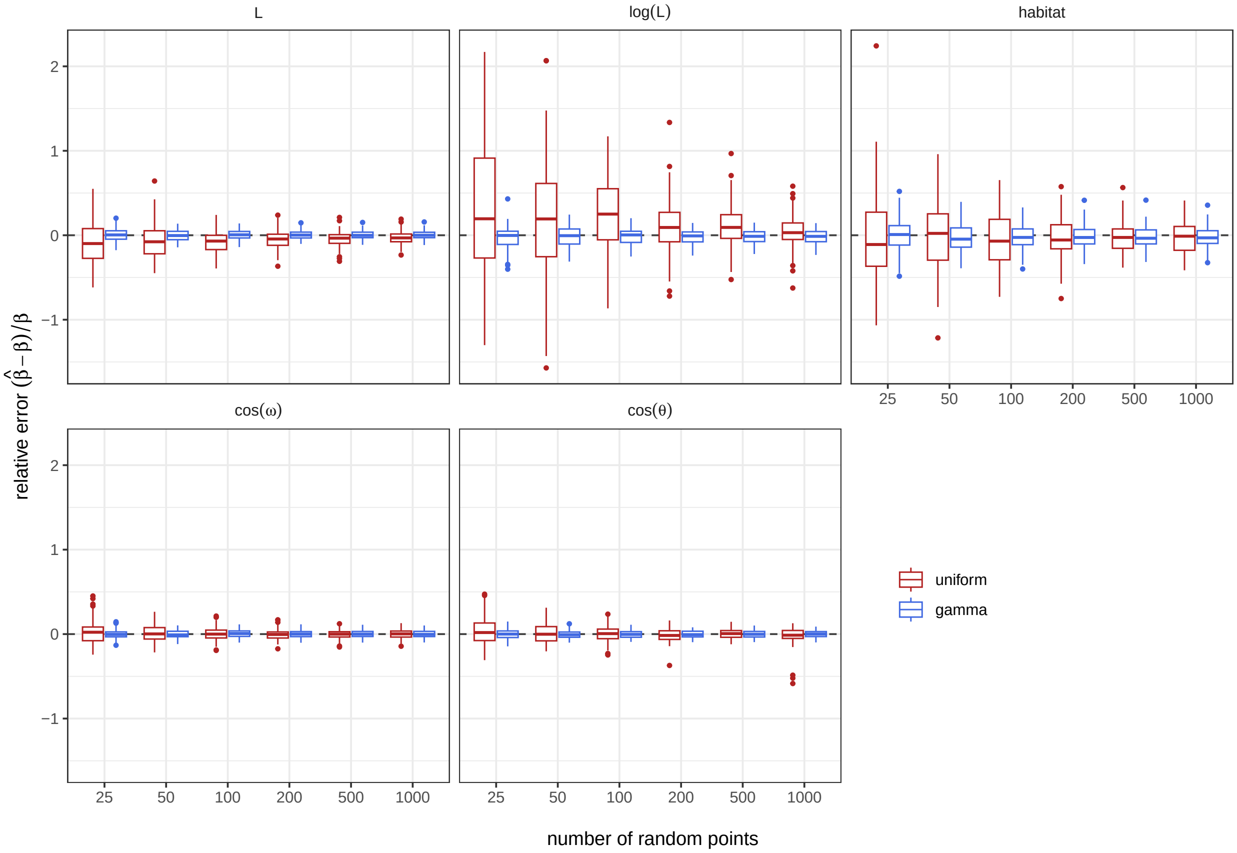

***Figure D3:*** *Results of the simulation study to check SSF implementation for the BCRW, with attraction towards level bearings. Note that the coefficients for* $\cos(\omega)$ *and* $\cos(\theta)$ *correspond to* $\kappa_1$ *and* $\kappa_2$ *directly.*

### E Additional details of petrel analysis

#### E.1 Data processing

**Location data** The temporal resolution varied between individuals and was irregular within individuals. Therefore, we only retained data from individuals where the median time interval was $\leq$ 30 minutes, and split tracks where there was a gap larger than 35 minutes. Most individuals had approximately a 5, 10, or 15 minute resolution. Therefore, we regularised the two-dimensional locations to a 10-minute resolution for all individuals using a continuous-time correlated random walk, implemented in the `crawlWrap` function of `momentuHMM` (Johnson et al., 2008; McClintock and Michelot, 2018). Then, we used linear interpolation to obtain an altitude for each regularised location. Although linear interpolation is a relatively simplistic form of regularisation, we only allowed very small gaps such that we were only ever interpolating across a maximum of 3 observations. We also defined whether each track segment was a departure or return, based on whether the petrel moved away or towards the colony. Note that the publicly available locations were already truncated to remove locations over water, and so each track segment could only be part of one of those trip phases (i.e., foraging at sea was removed).

**Wind data** We obtained three-dimensional ERA5 hourly wind data (web link, will add citation later). The data consisted of wind vectors at 14 pressure levels, and geopotential $h$. The pressure levels were converted to meters above sea level (i.e., altitude; $A$) following

$$A = \frac{R(h/G)}{R - (h/G)}, \tag{30}$$

where $R$ is the the radius of the Earth (assumed to be 6378000m), and G is Earth's gravitational acceleration (9.80665 $\text{ms}^{-1}$) (**?**). We interpolated wind vectors to petrel locations by finding the two closest altitude layers (i.e., below/above), interpolating the vectors of each layer to the two-dimensional locations, and then linearly interpolating between the two layers.
